# Selection inference in a complex genomic landscape: the impact of polymorphic inversions

**DOI:** 10.1101/2025.07.01.662526

**Authors:** Julia Morales-García, Wiesław Babik, Piotr Zieliński, Krystyna Nadachowska-Brzyska

## Abstract

Understanding the genetic basis of adaptation is a key objective in evolutionary biology. Although advances in genomic selection scans have greatly improved our ability to detect signatures of adaptation, distinguishing true signals from false positives remains challenging. This task is particularly difficult in regions of reduced recombination, such as polymorphic inversions. In this study, we examine the genome-wide landscape of selection in the European spruce bark beetle (*Ips typographus*), Europe’s most destructive forest pest, which has one of the most complex inversion-associated recombination landscapes known. Using simulation-based analyses and whole-genome resequencing data from 312 individuals across 23 populations, we applied two complementary selection scan methods (nSL and Λ) to assess how inversions influence the detection of adaptive signals. Simulations revealed that the two selection scan methods differ in their susceptibility to false-positive signals within inversions and that partitioning data by inversion genotype (i.e., homozygote classes) can substantially reduce these errors. Consistent with the results of our simulations, our empirical data showed that inversions are highly enriched for selection signals when all inversion genotypes are analyzed but are depleted when only homozygotes are considered. This demonstrates that focusing on homozygote genotypes allows for a more precise identification of putative selection targets and haplotype-specific signals, as it overcomes the confounding effects of suppressed recombination in heterokaryotypic individuals. In contrast, the detection of selection in collinear regions was largely unaffected by the presence of inversions. Our findings highlight that inversions may play an important role in shaping adaptation, underscoring the need to account for species-specific genomic architecture when interpreting signals from selection scans.

**Author summary:** Detecting the genetic basis of adaptation is one of the main goals in evolutionary biology. However, it is often difficult to tell whether selection signals identified by genome scans truly reflect natural selection signatures or are simply artefacts of genome structure. In this study, we examined how polymorphic inversions affect the detection of selection in the European spruce bark beetle, one of Europe’s most destructive forest pests. First, we assessed two selection scan methods using simulation approach. Second, we tested the influence of polymorphic inversion on the selection inference using genome-wide data from 23 beetle populations. We showed that inversions can create false signals of selection when all individuals are analyzed together but by separating individuals according to their inversion genotypes, we were able to distinguish potential adaptive regions from false positives. Our results demonstrate that accounting for genomic architecture is crucial for identifying selection signatures and provide practical guidelines for studying species with inversion-rich genomes.

## Introduction

Elucidating the genetic basis of adaptation is one of the major goals of evolutionary biology [1]. The long-standing debate over whether selection acts primarily on many loci of small effect or few loci of large effect, and whether de novo mutations or standing genetic variation are more important in adaptation [1–8] has led to a large body of empirical adaptation research [1,9–10]. The progress in this field has been facilitated by a growing number of increasingly sophisticated methods for genomic selection scans [11–13]. Despite many methodological advances and the fact that selection leaves characteristic signatures around its targets, there are still serious limitations in distinguishing true selection signals from false positives. In particular, we still have a limited understanding of how regions of reduced recombination affect our ability to identify true selection signals and the general importance of such regions in adaptation [14–16].

Local reduction of recombination is one of the main characteristics of polymorphic inversions - chromosomal mutations that reverse the order of the sequence in the affected region and suppress recombination between two inversion haplotypes in heterozygous individuals. The effect of suppressed recombination may extend beyond the inverted region itself [16–20]. Furthermore, the recombination landscape in these regions is shaped by the frequency of inversion haplotypes - the more balanced the frequencies, the higher the number of heterozygous individuals in the population and the lower the population recombination rate in the inverted region [21]. A direct consequence of suppressed recombination between alternative inversion haplotypes is independent accumulation of mutations on both arrangements. This creates a genetic subdivision within the inversion region - a situation that is similar to a demographic scenario of two isolated populations [22–25]. In addition, gene flux (double-crossover and conversion events) that can happen between inversion haplotypes mimics a scenario of two isolated populations with some migration between them [20,23,26]. These inherent features of inversion polymorphism are important since free recombination (between loci) and panmixia are common assumptions of selection scans and many other inference methods. While new methods are often evaluated for their robustness to recombination rate variation and population subdivision [12,14,27], explicit inversion polymorphic scenarios have rarely been evaluated (but see [28] or [15]).

There are several reasons why including inversion regions in selection inference is important. First, theoretical investigations clearly showed that selection plays a key role in the establishment, spread and the long-term fate of inversions [16,29–35] and unlike other mutations, inversions are rarely selectively neutral [31,36]. Polymorphic inversions can be maintained by divergent selection and migration (e.g. inversion captures locally adapted alleles that are kept together in the face of gene flow [33]), different types of balancing selection [37–38] or, most likely, by a combination of these and other evolutionary processes (for a review, see [36,39]). Importantly, the relative roles of the different selection forces, drift and new mutations that accumulate within different inversion haplotypes (including both advantageous and deleterious ones) may change over time. Second, there is a growing body of empirical research that identifies polymorphic inversions as key drivers of adaptation that capture genes underlying traits involved in local adaptation [40–46]. Finally, accumulating evidence suggests that genomes harboring multiple polymorphic inversions and characterized by complex recombination landscapes may be more common than previously thought (so-called inversion-rich genomes [40,47]). This provides an opportunity to investigate selection signatures among many inversions in a single genome.

Due to the dynamic nature of inversions, it can be difficult to detect and/or distinguish true selection signatures from false positive signals. However, simulation-based studies have demonstrated that adaptive inversions carrying locally adaptive loci can be distinguished from neutral ones [16]. Yet it has also been shown that several selection scan methods can produce strong false positive signals within inversions [15]. Studies have also shown that inversions not only capture beneficial alleles but can also accumulate new advantageous mutations over time [16]. These mutations can undergo selective sweeps within inversion homozygotes and should be detectable in datasets containing the relevant haplotype, unless the underlying phenotypes are polygenic. In this case, the usual challenges of identifying selection signals at loci of small effect apply [2,15–16,48–49].

Here, we performed both simulation-based and empirical assessments of two selection scan methods using the inversion-rich genome of the spruce bark beetle [50] and provide guidance for methods usage in such genomes. The evaluated methods were the number of segregating sites by length (nSL) [27] and Λ from the saltiLASSI maximum likelihood approach [12]. We focused on methods that do not require recombination rate information, as this is still lacking for many of the species for which genomic data are being generated. Both methods are designed to search for signatures of hard and soft sweeps of ongoing or recent selection, but they differ in how they are affected by reduced recombination. While nSL has been shown to be quite robust to variation in recombination rate, Λ tends to be negatively correlated with recombination rate and strong Λ signals in low-recombination regions must be interpreted with extra caution [12,27]. Guided by simulation results we tested if applying selection scan methods in datasets restricted to inversion homozygotes reveal haplotype specific selection signals. Finally, we identified candidate loci under selection within the spruce bark beetle genome.

## Materials and Methods

### Study system

The European spruce bark beetle (*Ips typographus*, hereafter the spruce bark beetle) is found in Norway spruce forests throughout Eurasia. It is often considered to be the most destructive forest pest, which under certain conditions can cause mass mortalities of spruce stands [51–52]. In recent decades, exacerbated by climate change, the frequency and severity of outbreaks have dramatically increased, making the spruce bark beetle a major economic and environmental concern in European forests [53–55]. Extensive research on species ecology and biology, as well as, on causes and consequences of the spruce bark beetle outbreaks, has been recently complemented by population genomic studies [50,56]. Importantly, these studies identified 27 large polymorphic inversions covering approximately 30% of the species genome [50]. Most of the identified inversions are polymorphic across species range but can significantly vary in frequency among populations which makes the spruce bark beetle an excellent model to study natural selection signatures in the context of inversion rich genome.

### Sampling, sequencing, and data preparation

We used data presented in Mykhailenko et al. (2024) [50] and Zieliński et al. (2025) [56]. The data included 312 individuals from 23 populations (13-14 individuals per population). Samples covered most of the species’ range in Europe and ranged from northern Italy to northern Fennoscandia (Figure 1; Table S1). The details of library preparation, sequencing and processing are given in Mykhailenko et al. (2024) [50] and Zieliński et al. (2025) [56]. In short, DNA was extracted from the whole beetles using the Wizard Genomic DNA Purification Kit (Promega). Genomic libraries were subject to 2×150 bp paired-end sequencing using the NovaSeq technology (Illumina Inc.). Raw data was subject to quality control with FastQC [57]. Low quality reads and adaptors were trimmed with Trimmomatic [58] and mapped to the spruce bark beetle genome [59] with Bowtie2 [60]. Duplicate reads were removed with PicardMarkDulicates (https://broadinstitute.github.io/picard/). Variant calling, genotyping, base quality score recalibration, and filtering were performed with GATK, following the GATK best practice guideline. Filtering included: removing five sites around indels and sites within repeat-masked regions of the genome, and variants exhibiting: low-quality (mapping quality below 30), total depth higher than mean + 1SD, excessive heterozygosity. Individual genotypes with coverage lower than 8 and genotype quality lower than 20 were treated as missing. In downstream analysis we used only biallelic SNPs with more than 50% of genotypes called. The data for all individuals was phased using default parameters of Beagle 5.2 [61]. All analyses were performed using contigs larger than 1 >Mb (IpsContigs 1-36, 78% of assembly length). Moreover, IpsContig33 which had high similarity to mtDNA and potentially sex linked contigs were excluded (IpsContig9, IpsContig27, IpsContig31 and IpsContig35, Nadachowska-Brzyska in preparation).

**Figure 1.**
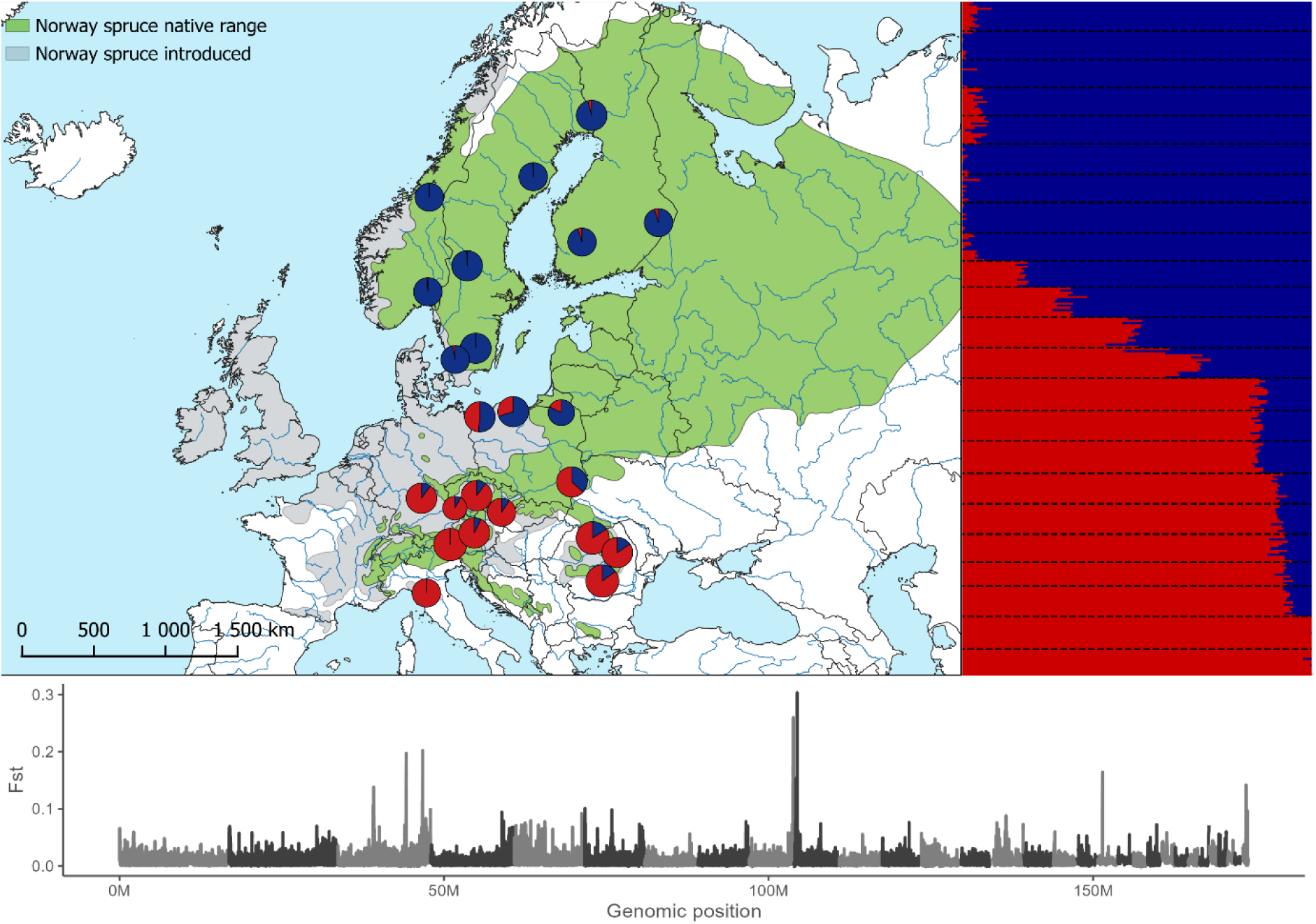
The geographic distribution, admixture, and genetic differentiation of *Ips typographus* populations as inferred in Zielinski et al. 2025. Top left: map of sampled populations, with admixture proportions illustrated by coloured circles. The colours represent the admixture proportions that were identified (top right). Admixture plot showing individuals admixture proportions for K=2. The populations are separated by dashed lines and ordered according to longitude. Bottom plot: pairwise Fst calculated between the two genetic clusters across the genome. Grey and black represent different contigs.

### Selection scans

nSL was calculated in selscan 2.0.2 [62] using default parameters and phased data (it is not possible to use unphased data if it contains missing genotypes - a very common situation with whole-genome resequencing data). The nSL values for each SNP were normalised against genome-wide distribution using ‘norm’ function and default settings. Because the sign of nSL depends on the unknown ancestral versus derived state of the allele, we followed standard practice and used the absolute values to capture both types of signatures. The top 1% of nSL absolute values were treated as outliers. Λ was inferred using LASSI-Plus1.1.2 [12]. In contrast to the nSL approach, which provides statistics for each analysed SNP, Λ is given for windows consisting of a predefined number of SNPs (hereafter called SNP-based window). In addition, different window sizes may be more suitable for detecting recent or more ancient sweeps [12].

We tested different window sizes and based on our tests and previous studies [12], we decided to use two window sizes: 52 (sliding window step 12) and 117 SNPs (sliding window step 12). The top 1% of Λ values were considered outliers.

The nSL results based on the spruce bark beetle data were translated into SNP-based Λ windows and mean values of nSL were calculated in each window. This was done to be able to statistically compare selection signals between statistics. The top 1% mean (window) nSL absolute values were considered outliers. In order to be able to compare different datasets that contained different individuals and thus different numbers of SNPs and different numbers of SNP-based windows, we further standardised the results by summarising the statistics (both nSL and Λ) in non-overlapping 50 kb genomic windows. For each 50 kb window, we calculated the mean value of each statistic of all SNP-based windows that overlapped it. This allowed a consistent window-based comparison across datasets and statistics.

Finally, to test if selection signals tend to cluster within inversions, we plotted outlier windows (SNP or 50kb windows according to each case) along the genome, investigated the position of each outlier window in comparison to inversion coordinates and calculated the sizes of clusters composed of adjacent outlier windows. Additionally, to correlate selection statistics with the genetic differentiation between inversion haplotypes we summarised Fst in SNP-based windows (in the same way as we did for nSL results) using vcftools [63].

### Simulations

To evaluate the effect of inversions on selection scan methods we first investigated selection scans performance using simulations (Figure 2a). We based our test on the existing and publicly available set of simulation performed by Lotterhos (2019) [15] who created a resource to test new methods and their sensitivity to realistic patterns of genomic variation, including polymorphic inversion. In short, the author performed 200 replicate forward-time simulations of a metapopulation adapting to a heterogenous spatial environment. The simulations were performed in SLiM v 3.2 [64]. The simulated data included nine linkage groups (LGs) each evolving independently and under different evolutionary scenarios: two LGs followed neutral evolution (LG1 and LG2), and the rest included multiple casual quantitative trait nucleotides (LG3 and LG4), complete selective sweep (LG5), partial sweep (LG6), inversion maintained polymorphic by weak frequency dependent selection (LG7), low recombination region mimicking centromere (LG8) and randomly varying recombination across the LG (LG9). Each LG was 50kb and 50cM in length and simulations included 1000 individuals adapting to heterogenous spatial environment. Simulated inversions were 10kb long and were located in the middle of the LG7, their fitness was not related to local adaptation to the environment and all mutations within the inversion were neutral. For our evaluation we used vcf files deposited at Dryad ([15] results from 181 replicate simulations). In each simulation, individuals were classified according to their inversion genotypes using the approach described in Mykhailenko et al. (2024) [50]. Reliable inversion genotyping was possible for 179 of the 181 simulations. Classification accuracy was verified with inversion PCA plots, which showed individuals clustering distinctly into three genotype groups, and were coloured according to the inferred inversion genotypes (Supplementary Information: Simulations Results .zip folder).

**Figure 2.**
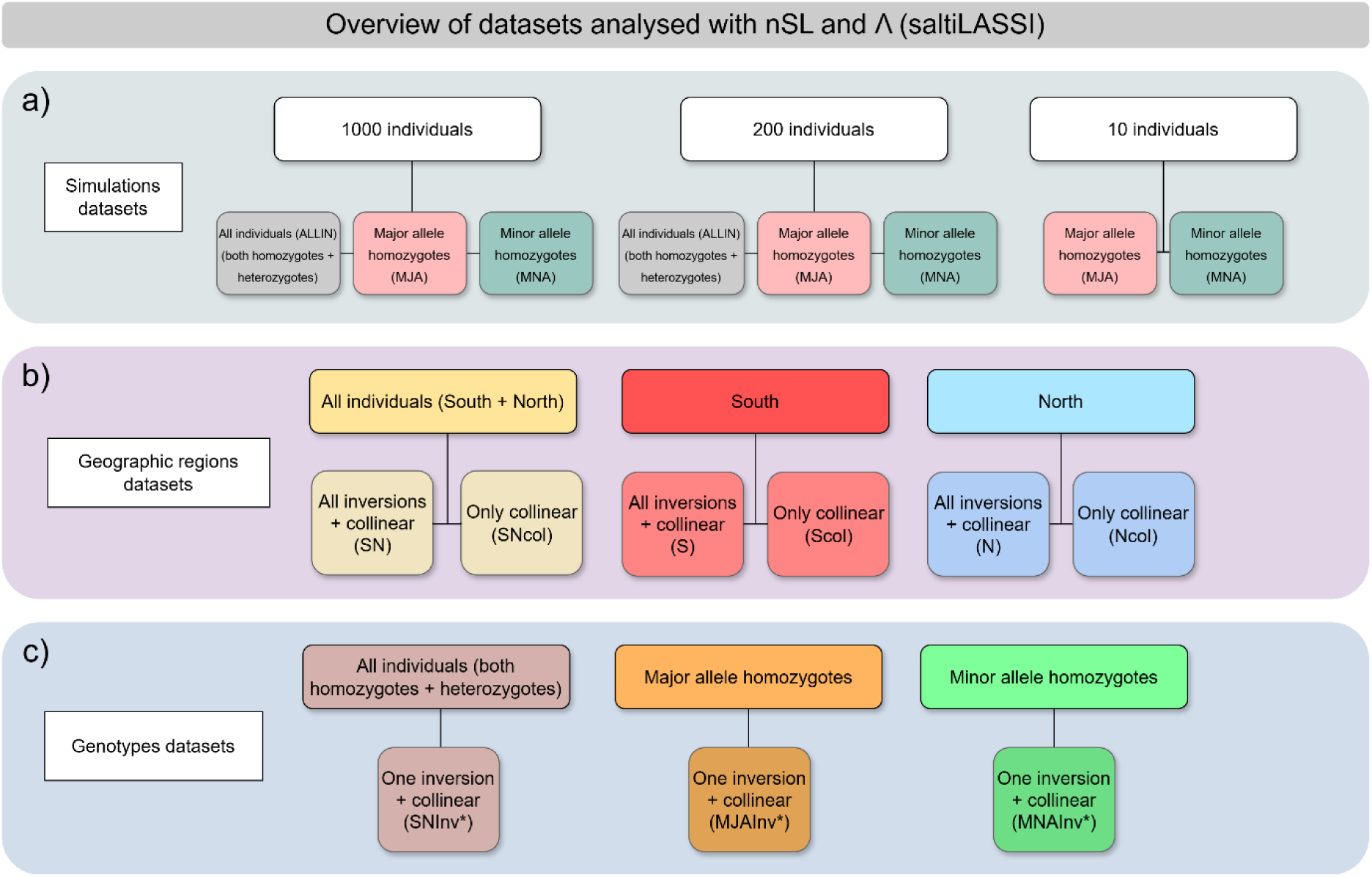
Graphical summary of the datasets analysed with nSL and Λ selection scans. A: Simulation datasets prepared based on the publicly available simulation results from Lotterhos (2019). The datasets differ in samples size (1000, 200, and 10 individuals) and included all individuals (all possible inversion genotypes, ALLIN, grey), or only individuals homozygous for a particular inversion allele (major allele; MJA, pink and minor allele, MNA, turquoise). B: Geographic region datasets: datasets generated from the spruce bark beetle genomic data and including southern populations (S, red), northern populations (N, blue) or both combined (SN, yellow). Each group was analysed including all genomic regions (inversions + collinear) or restricted to analysing collinear regions only (Scol, Ncol and SNcol). C: Genotype-based datasets: generated from the spruce bark beetle genomic data and analysed either including all individuals, regardless of their inversion genotype (both homozygotes and heterozygotes; SNInv*, brown), or including only individuals homozygous for a particular inversion allele, either for the major (MJAInv*, red) or minor (MNAInv*, turquoise) allele. Each inversion was examined separately, and generated dataset consisted of the inversion in question and collinear part of the genome. Inv* indicates 24 inversions examined.

Selection scans were conducted using all replicate simulations and all individuals (including all inversion genotypes), as well as datasets consisting of homozygotes of one type only (Figure 2a). Additionally, all simulations were downsampled from 1000 to 200 individuals (similar sample size as the spruce bark beetle sample size) to evaluate the impact of sample size on selection statistics. Since homozygote datasets in the spruce bark beetle data often substantially differ in sample size, in particular if one inversion haplotype is rare, we also downsampled homozygote datasets in simulated data to 10 individuals. We used the selection scan results across these datasets to test whether the two evaluated methods of selection scans produced false positive signals within neutral inversions and whether sample size affected the inferred selection landscape within inversions. To assess false positives, we counted the number of outliers within inversion regions and compared this to the number of outliers detected in neutral linkage groups (LG1 and LG2; chi-squared tests were used to compare the proportion of outliers in inversion versus collinear regions). This test was conducted for datasets including all individuals, homozygote-only datasets, and all replicate simulations. To examine the effect of sample size, we used generalized linear mixed models (GLMM; *glmmTMB* package [65]; Supplementary Information: GLMM.R) with a binomial response representing the number of outlier SNPs and non-outlier SNPs detected in inversion region. The simulation replicate was included as a random effect to account for the variation between simulations. The fixed effects were sample size (200 versus 1000 individuals in datasets including all genotypes, and 200 versus 10 individuals in datasets including only homozygotes). Through this approach, we investigated whether a sample size of approximately 200 individuals (comparable to the spruce bark beetle dataset) would yield selection scan outcomes within the inversion region that mirrored those achieved with a sample size of 1000 individuals. We also tested whether a very small sample size of 10 individuals would produce unbiased results similar to those obtained with a dataset of 1000 individuals divided into homozygotes.

### Datasets, tested hypotheses and statistical tests

Multiple spruce bark beetle datasets used in the study are presented in Figure 2b and 2c. Individuals were grouped according to the previously described genetic structuring into the southern and northern clade ([50,56]; Figure 1, Table S1) creating three datasets: 1) all individuals from southern and northern region (SN, 258 individuals), 2) only individuals from the southern populations (S, 138 individuals), 3) only individuals from the northern populations (N, 120 individuals). These datasets did not include Polish populations that are highly admixed. Each of these datasets was analysed in two versions, differing in the included parts of the genome: inversion regions were either included (SN, S, N) or excluded (SNcol, Scol, Ncol; only included collinear genome).

To investigate the effects of selection in inverted homozygotes, all individuals were also divided into datasets based on their inversion genotype, forming three datasets per inversion: 1) individuals carrying major inversion haplotype (e.g. MJAInv2); 2) individuals carrying minor inversion haplotype (e.g. MNAInv2); 3) individuals carrying all possible inversion genotypes (e.g. SNInv2). Polish individuals were included in this analysis to ensure sufficient sample sizes, and because this analysis focused on differences between inversion genotypes rather than population-level structure. Inversion datasets consisted of inversion in question and collinear part of the genome (thus, excluding other inversions apart from the one in question).

### Datasets with and without inversions

The analysis of the datasets with and without inversions (SN, S and N, and their respective SNcol, Scol and Ncol, Figure 2b) allowed us to investigate whether all or some inversions appear as selection outliers, to test whether the presence of multiple inversions affects the detection of selection signals outside of inversions (in collinear part of the genome), and to compare the behaviour of the two methods in these respects. Specifically, we predicted: 1) to find more selection signals in inversions than in collinear parts due to the effect of reduced recombination and inversions’ genetic subdivision, 2) that balanced inversions with intermediate frequency of both inversion haplotypes will show stronger selection signals than inversions with one allele rarer (assuming that rare allele is younger and have low within haplotype polymorphism), 3) that outlier detection outside inversions is affected by inversions present within the genome due to their influence on haplotype frequency spectra (Λ) and the assignment of the statistics values into the frequency bins (nSL). To test whether selection signals are more often found in inversion regions than in collinear parts of the genome we calculated the proportion of outliers in inversions and performed a permutation test for each of the analysed dataset (1000 permutations, permuting SNP-based window labels - inverted and collinear). To check the effect of inversion polymorphism frequency on selection scan performance, we correlated the inversion haplotype frequency and the percentage of outliers within the inversion. To test whether inversions affect selection inference in the collinear part of the genome we compared datasets with and without inversions. We counted the number of outlier windows and investigated whether the same regions were classified as outliers in both datasets and whether new selection signals appear if we excluded inversions from the analysis. We compared both methods by checking the number and distribution along the genome of the outlier windows identified in both methods and across all datasets.

To be able to identify selection signals specific for the genetically differentiated groups identified across the species range, selection scans described above were performed separately for southern and northern regions/clades (S and N; Scol and Ncol; Figure 1, Figure 2b). To test if selection signals correlate with Fst between the southern and northern group we correlated Fst with nSL and Λ values (all calculated in SNP-based windows, significance of the correlation was calculated using permutations).

### Datasets consisted of different inversion genotypes

The inversion datasets (MJAInv, MNAInv and SNInv, Figure 2c) allowed us to test whether: 1) selection signals present in inversions when analysing dataset with all individuals (SNInv; all possible inversion genotypes) disappear when the selection scans were performed using only homozygous individuals (MJAInv or MNAInv); 2) There is a positive association between inversion polymorphism and signal disappearance (more false positive selection signals expected in high frequency of heterozygotes); 3) Selection signals disappear only in one homozygotes revealing haplotype which carries alleles under selection. We expect that the behaviour of the scans depends on the frequency of the inversion haplotype (produced by lower recombination within haplotypes and higher genetic subdivision between them) and that the most pronounced changes in selection signal occur when the inversion frequency is the highest (around 0.5). We also predict that in some inversions false positive signals would disappear, leaving only inversion haplotype-specific selection signals (if they exist).

In total, twenty-four inversions were analysed. In each of the inversion dataset all the other inversions (apart from the inversion in question) were excluded before performing selection scans. We calculated the percentage of outliers in each inversion in all datasets and in both of the selection scans. Since 1% of the selection scans results are always treated as outliers (even if the selection landscape is flat) we corrected the observed percentage of outliers in each inversion for the expected number of outliers given the inversion length and analysed genome size (given in number of SNP windows). Since several observed percentages equalled zero, we did not normalize them by simply dividing observed by expected values. Instead, we transformed the observed data as (observed percentage - expected percentage)/(observed percentage + expected percentage)/2. This produced values that were easy to interpret: positive values indicated regions enriched in the selection signal and negative values regions that were depleted in the selection signals. These transformed percentages were used in all the downstream tests.

To test for a difference between the observed and the expected percentage of outliers within inversions and to test whether inversions are enriched in (Observed – Expected > 0) or depleted (Observed – Expected < 0) from selection signals we used Wilcoxon signed-rank test. To test whether the magnitude of selection signals depends on the frequency of the inversion haplotype, we fitted logistic regression models for each dataset (SNInv, MJAInv, MNAInv). We transformed the percentage of outliers into binary categories—“Depleted” (negative values) and “Enriched” (positive values). These categories were used as response variables, and the frequency of the major allele and its square (to capture a potential nonlinear relationships) were included as explanatory variables. Finally, to test if selection signals disappear only in one homozygote (major or minor) revealing haplotype which carries loci under selection we classified selection signals into two signal categories: 1) selection signal gets much stronger in one of the homozygous dataset (percentage of outliers in one MJAInv (or MNAInv) dataset is higher than 50% of the percentage observed in the SNInv dataset (all genotypes) and the percentage of outliers in the other homozygote dataset is lower than 50% of the percentage observed in the SNInv dataset); 2) percentage of outliers is different from the first category. After categorization we used binomial regression model with signal behaviour as response variable and frequency of the inversion major haplotype as explanatory variable. The significance of the variables was tested using a Wald test.

### Candidate genes under selection and gene ontology enrichment analysis

We identified genes harbouring or closest to outlier windows from the top 1% of nSL or Λ values. We used gene annotations from Powell at. al (2021) [59] and tested for overrepresentation of GO using R package topGO [66]. We applied Fisher’s exact test and “weight01” algorithm [67] to deal with the GO graph structure and only GO categories with at least 10 members among the SNP associated genes were considered. GO analysis were performed for datasets including: 1) selection outliers outside inversions and specific for northern group; 2) selection outliers outside inversions and specific for southern group; 3) selection outliers outside inversions common for both groups; 4) selection outliers specific for particular inversion haplotype or present in both inversion haplotypes (the inversion signals that we consider true signals).

## Results

First, we used simulations to assess the performance of the nSL and Λ selection scans and to establish a theoretical baseline for the patterns expected in neutral inversions across the genome. Second, we used whole-genome sequence data from hundreds of individuals and performed both selection scans across the inversion-rich spruce bark beetle genome. The list of all datasets and their characteristics (e.g. whether they contained inversions or not) are given in Figure 2 and the results are presented below.

### Selection landscape in neutral inversions - lessons from simulation-based datasets

#### 1. False positive signals in neutral inversions

Neutral inversions contained selection outliers in the majority of simulations (Figure 3, Supplementary Information: Simulations Results .zip folder). This pattern was consistent across datasets with 1000 and 200 individuals and for both selection scans (nSL and Λ, Table 1).

**Figure 3.**
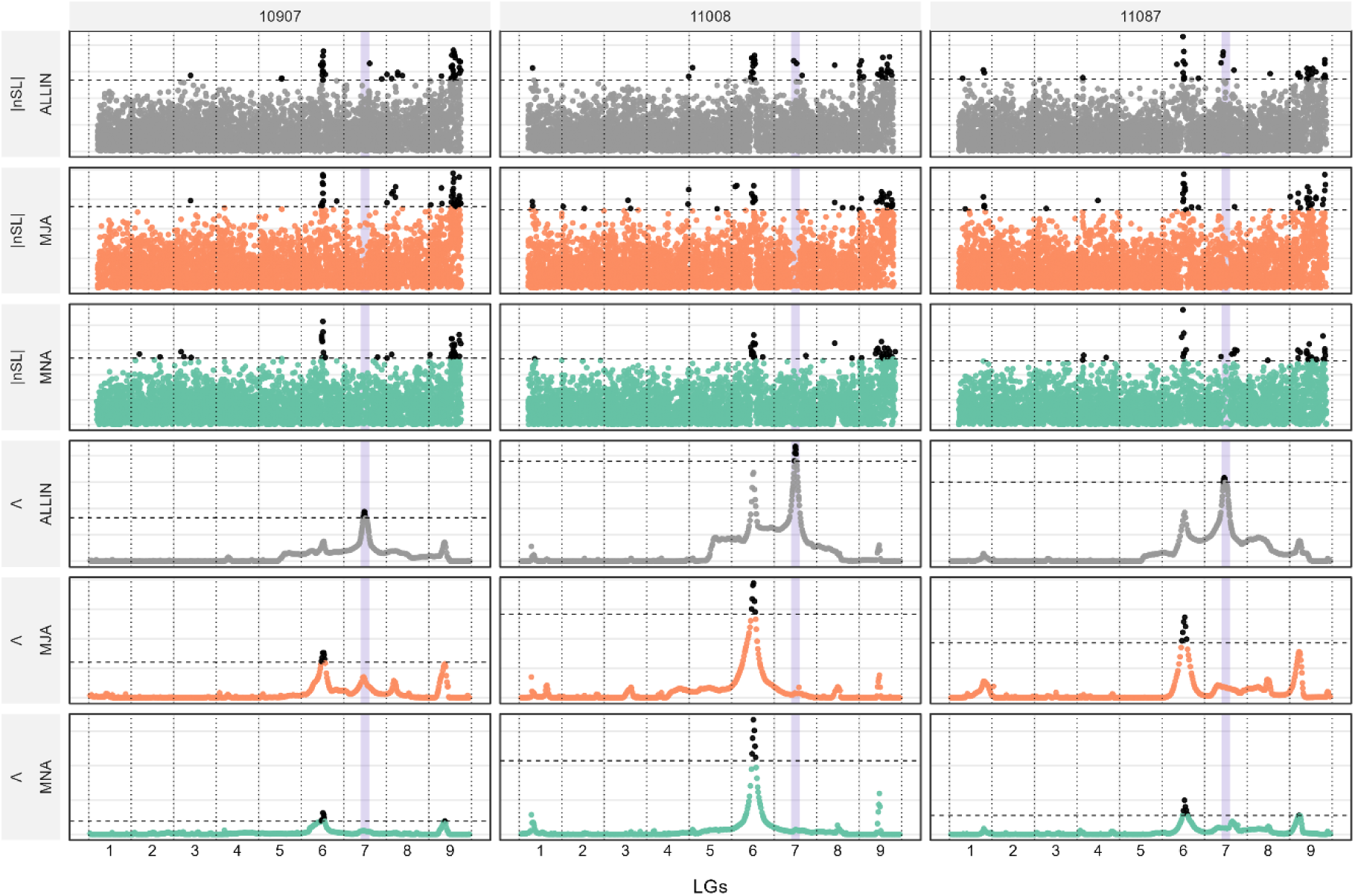
Comparison of selection scan results in three example simulations. Each column represents an independent simulation (numbered according to Lotterhos 2019), and each row corresponds to a dataset and selection statistic. The rows show either the nSL or the Λ statistic, which were computed for three datasets based on the inversion genotype of individuals: ALLIN (both homozygotes and heterozygotes, grey); MJA (major allele homozygotes, orange); and MNA (minor allele homozygotes, green). The dashed horizontal line indicates the 99th percentile threshold and the black dots above this line represent selection outliers. The vertical dotted lines mark the boundaries between linkage groups (LGs), which are numbered along the x-axis. The purple shaded region highlights the inversion region. LG1 and LG2 evolve neutrally and the selection signals within them were compared to selection signals detected within inversion. The most pronounced selection outliers were detected in LG6 included partial selective sweep.

**Table 1.** Percentage of simulated datasets in which at least one outlier was detected within the inversion. Calculations were performed for datasets with 1000 and 200 individuals and for both Λ and nSL selection scans. Within each dataset, individuals were grouped according to their inversion genotype into three categories: ALLIN: all individuals, including both heterozygotes and homozygotes; MJA: including only individuals homozygous for the major inversion haplotype; MNA: including only individuals homozygous for the minor inversion haplotype.

|  |  | 1000 individuals |  | 200 individuals |  |
| --- | --- | --- | --- | --- | --- |
| | | nSL | $\Lambda$ | nSL | $\Lambda$ |
| % of simulations with at least one outlier in the inversion | ALLIN | 84% | 59% | 83% | 60% |
|  | MJA | 32% | 0% | 32% | 2% |
|  | MNA | 54% | 1% | 41% | 1% |

The two selection scan methods revealed qualitatively different patterns of selection signals. When present, Λ outliers produced distinct and pronounced peaks, with no outliers detected in the two linkage groups that evolved under neutrality in the simulations (Figure 3 and Supplementary Information: Simulation Results .zip folder). In contrast, nSL scans typically identified only a small number of outlier SNPs within inversions (Figure 3 and Supplementary Information: Simulation Results .zip folder). This pattern was very similar to patterns observed in neutral LGs (LG1 and LG2) where a small number of outliers were also detected. In most simulations, the proportion of outliers in inversions and neutral linkage groups did not differ significantly (χ² test, *p* > 0.05 after multiple testing correction). In the dataset with 1000 individuals, 33 simulations showed significantly more outliers in inversions compared to neutral linkage groups, while in the dataset with 200 individuals, this was the case for 31 simulations.

#### 2. What happens to section signals if we divide simulations into homozygotes datasets?

A striking difference emerged between the two selection scan methods in how selection signals changed when datasets were divided into major and minor inversion homozygotes and compared to the simulations that showed selection signal in datasets that included all inversion genotypes. In Λ scans, nearly all previously detected signals disappeared in the homozygote subsets (Figure 3 and Supplementary Information: Simulations Results .zip folder, Table 1). In contrast, nSL scans showed a more stochastic pattern of signal loss and appearance (Table 1). As in the previous analyses, only a small number of simulations showed a significant excess of outliers in inversions compared to neutral linkage groups (χ² test, *p* < 0.05 after multiple testing correction), with between 2 and 10 out of 181 simulations showing a significant difference.

#### 3. The effect of dataset sample size on the detection of selection signals

We found no significant difference in the proportion of outliers in inversions between datasets of 200 and 1000 individuals for either nSL or Λ selection scans (GLMM, *p* > 0.05). In these comparisons, sample sizes were fixed at 200 and 1000. When datasets were divided into homozygote subsets, the sample size depended on inversion frequency, and even in the 1000 individual dataset, minor haplotype subsets could be as small as 10 individuals (3 simulations were excluded since had only 1 or 2 homozygotes; in total there were 38 simulations had fewer than 100 individuals). For nSL scans, no significant difference was observed between major inversion haplotype datasets from 1000 individuals (smallest *n* = 208) and those with only 10 individuals. For minor inversion homozygotes, slightly fewer outliers were detected in the smallest datasets (*n* = 10) compared to larger subsets (GLMM, *p* = 0.01).

### Selection landscape in inversion-rich genome – lessons from empirical datasets

#### 1. Inversion regions are enriched in selection signals

Most of the selection outliers were identified within inversion regions (Figure 4, Figure S1, Table 2), many more than expected by chance (permutation tests; p < 0.001; Supplementary Information 1.1). This pattern was consistent across both statistics, window sizes and all datasets (SN, S, N), except for Λ scan and SN dataset and 52 SNPs window size, where more outliers than expected were found in the collinear part of the genome (Table 2). In the latter case the signal was mainly driven by one region in IpsContig3 (39% of all outliers and 55% of collinear outliers in IpsContig3). Only 7 inversions (out of 24) did not show any selection signals in nSL or Λ analysis (Table S2). Inversions not only included more outliers in total but also harboured longer selection signal clusters (in particular in Λ scans; permutation test, p < 0.001; Table S3; Supplementary Information 1.8). This was also true for the Λ scan with 52 SNPs window size that had a smaller percentage of outliers in inversions compared to the collinear part of the genome, even considering that inversions represent only ∼30% of the genome (Figure 4, Table S3). The largest inversion outlier clusters were found in inversion Inv16 (0.324 Mb, 3% of the inversion, in S dataset) and Inv5 (0.253 Mb, 6% of the inversion, in N dataset, Figure 4, Table S3). The largest cluster outside inversion was found in IpsContig3 (ranging from 0.317 to 0.369 Mb in different datasets). We did not find a significant correlation between frequency of inversions and strength of selection signals (Supplementary Information 1.3). In summary, we found that inversions are enriched in selection signals and that these signals form large blocks of outlier regions.

**Figure 4.**
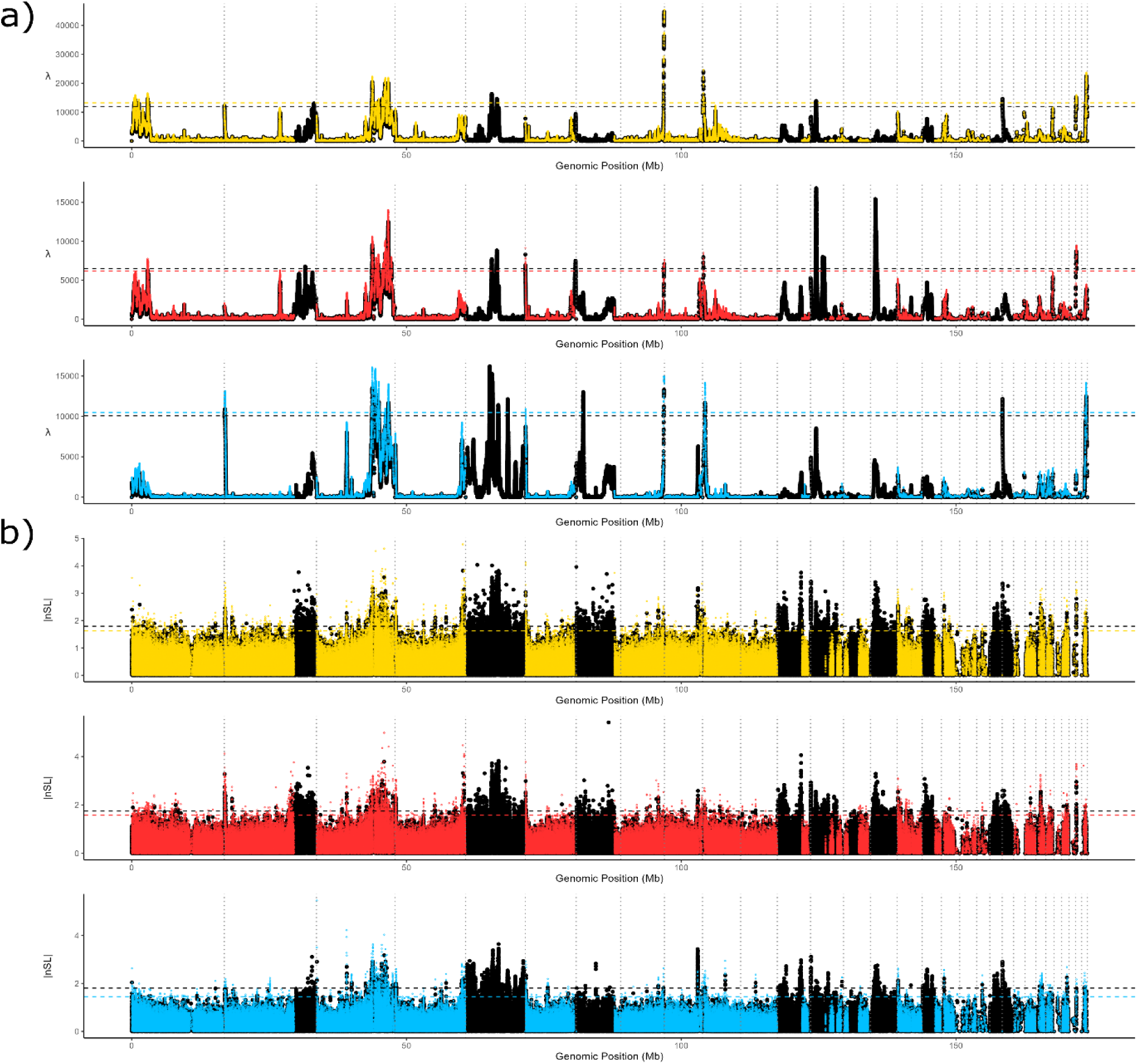
Results of the selection scans for the 52 SNP window sizes and SN, S, and N datasets for both selection scans (a) Λ and (b) nSL (absolute values). Results using both collinear and inversion regions are shown in black and the results of the same analysis performed excluding inversion regions are overlaid (SNcol – yellow, Scol – red, and Ncol – blue). The dashed lines indicate the outlier threshold (1% of all values). In all plots, grey dotted vertical lines represent the end of each contig.

**Table 2.** The comparison between Λ and nSL selection scans using different datasets and a 52 SNP window size. Calculations from nSL analysis are based on the absolute values of the statistic. Genomic region: part of the genome that is included in each dataset. Inversion-associated outliers: the percentage of outlier windows located within inversion regions, calculated from the total number of outliers. Collinear-region outliers: percentage of outliers located outside inversion regions calculated from the total number of outliers. Shared outliers: percentage of outliers that are shared between Λ and nSL results. Shared outliers in inversions: the percentage of outliers shared between Λ and nSL results that are in inversion regions. Shared outliers in collinear: the percentage of outliers shared between Λ and nSL results that are in the collinear genome.

| Dataset | Genomic region | Statistic | Inversion associated outliers | Collinear region outliers | Shared outliers | Shared outliers in inversions | Shared outliers in collinear |
| --- | --- | --- | --- | --- | --- | --- | --- |
| SN | Inversions + collinear | $\Lambda$ | 29% | 71% | 15% | 12% | 3% |
|  |  | nSL | 86% | 14% |  |  |  |
| SNcol | Only collinear | $\Lambda$ | | 100% | 11% | | 11% |
|  |  | nSL |  | 100% |  |  |  |
| S | Inversions + collinear | $\Lambda$ | 70% | 30% | 20% | 18% | 2% |
|  |  | nSL | 87% | 13% |  |  |  |
| Scol | Only collinear | $\Lambda$ | | 100% | 16% | | 16% |
|  |  | nSL |  | 100% |  |  |  |
| N | Inversions + collinear | $\Lambda$ | 56% | 44% | 17% | 15% | 2% |
|  |  | nSL | 91% | 9% |  |  |  |
| Ncol | Only collinear | $\Lambda$ | | 100% | 19% | | 19% |
|  |  | nSL |  | 100% |  |  |  |

#### 2. Selection landscape identified in collinear parts is minimally affected by inversions present in the genome

Both nSL and Λ scans conducted using the dataset with inversions, revealed multiple signals of selection outside inversion regions (Table 2-3, Figure 4-5, Figure S1-S2). The proportion of outliers detected depended on the dataset and selection scan method used. Figure 5 and Table 4 compare the selection landscape in the collinear part of the genome in datasets that included inversions when the selection scans were performed (SN, S, N) and datasets in which inversions were excluded before selection scan analysis (SNcol, Ncol, Scol). There was a significant correlation between the results obtained in the collinear part of the genome using datasets that included and excluded inversions (r^2^ ranged from 0.76-0.95, permutation test; p-value < 0.001; Supplementary Information 1.6). In summary, the general selection landscape in collinear genome was almost the same between datasets that included and excluded inversions indicating minimal influence of inversions on the selection scan results in the collinear part of the genome (Figure 5).

**Figure 5:**
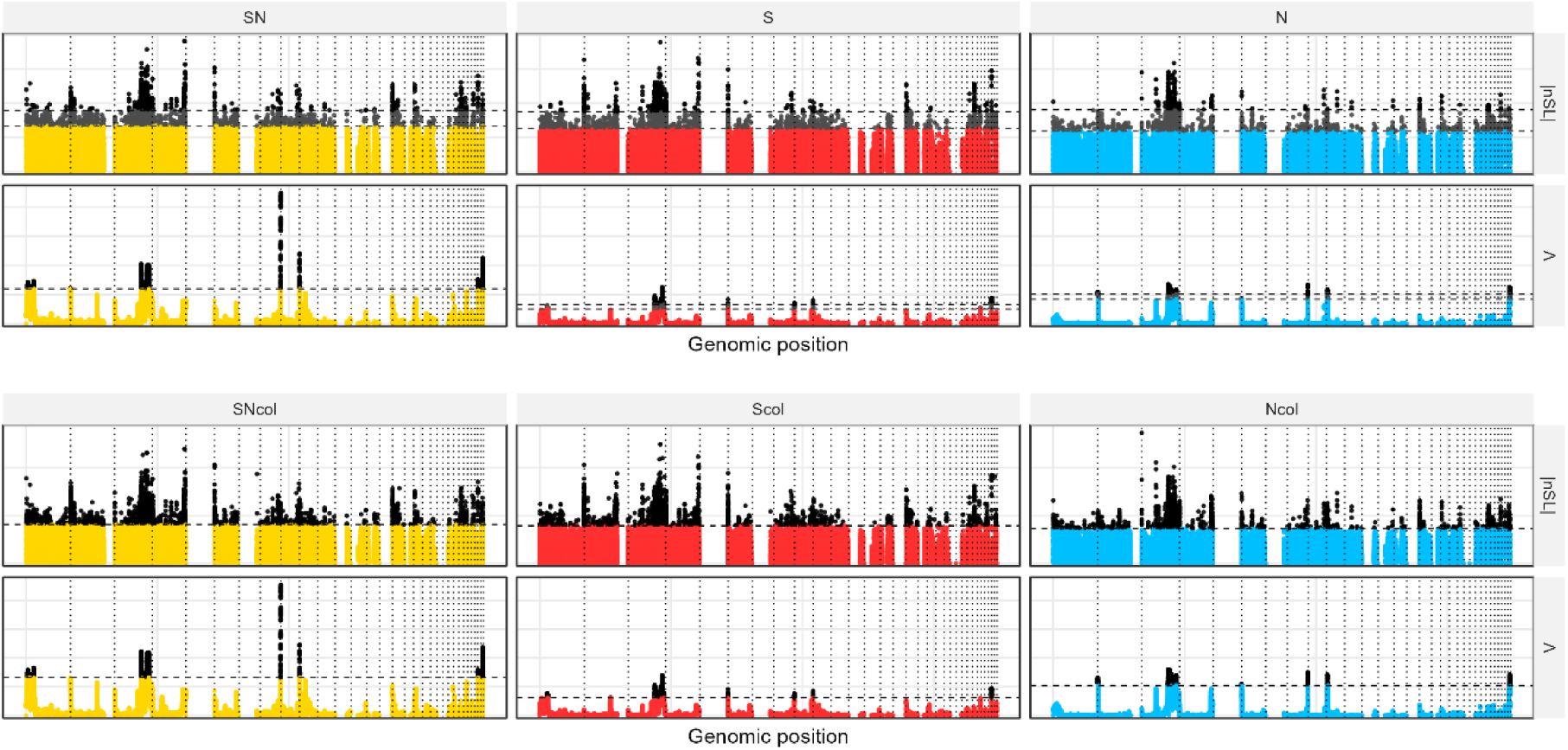
Results of the selection scans for 52 SNP window sizes and the SN, SNcol, S and Scol, N and Ncol datasets for Λ and nSL selection scans. Each column represents the results of the datasets relating to the geographic populations of the spruce bark beetle: SN (all individuals, south and north, yellow), S (south, red) and N (north, blue). Each row shows the results of the nSL or Λ analyses. The upper raw of plots shows the results of analyses performed with both collinear and inverted regions included, while the lower raw shows the results of analyses performed with only collinear regions. Every vertical dashed line differentiates a different contig. The horizontal dashed lines set the threshold at the 99th percentile, with outliers above shown in black and grey (depending on the threshold used). In the upper raw of plots, two horizontal dashed lines indicate the 99th percentile threshold: one was calculated including inversions (black outliers) and the other was calculated excluding them, with the corresponding outliers shown in dark grey and black. By comparing upper and lower raw one can see the inversions inclusion of inversions in the analyses does not affect the selection landscape in collinear genome.

#### 3. Both methods identify a similar selection landscape

Both selection scan methods produced a fairly similar selection landscape across all datasets (r^2^ ranged from 0.09 to 0.21, permutation test; p < 0.001; Supplementary Information 1.4). Only the selection landscape obtained from Λ scans with window size 117 SNPs was visually different and dominated by two large outlier clusters in inversions Inv5 and Inv7 (Figure S1, Table S4). Despite this difference, many regions identified in the analysis that used 52 SNPs window size (both nSL and Λ) were still visible as distinct peaks when using window size of 117 SNPs but were not classified as outliers in this dataset due to the extreme values of the two inversions mentioned above. In general, the genome-wide selection landscapes were similar across different statistics (as indicated by a high correlation of values in 50kb widows), but the actual overlap between outlier SNP-based windows in nSL and Λ was rather small due to the difference in values for particular windows - the same regions appear as outliers but particular small SNP windows can differ in the magnitude of selection signals (range from 11% to 20% of shared outliers in Λ and nSL for 52 SNPs window size in different datasets; Table 1), but this is still more overlap then expected by chance (permutation test, p < 0.001; Supplementary Information 1.4). In summary, both methods identify similar selection landscapes despite the observed difference in magnitude of selection signal between nSL and Λ.

#### 4. Northern and southern regions have similar selection landscapes with few region-specific selection signals

Selection landscapes inferred for S and N datasets were highly correlated with each other (r^2^ equalled 0.58 for Λ, 0.66 for nSL; p < 0.001; Figure 4, Figures S1–S2, Tables 2-3, Supplementary Information 1.5). This overall similarity was consistent across both nSL and Λ and reflected the largely shared genomic landscape of selection between geographic regions (including similar impact of inversions on selection landscape in both regions). However, a region-specific outlier windows were also detected (Figure S3, Table 3). In addition, both nSL and Λ values were very weakly positively correlated with Fst values calculated across the genome and between the N and S datasets (r^2^ ranged from 0.0004 to 0.0634 across datasets; p < 0.001; Supplementary Information 1.2).

**Table 3.** Comparison of the results of Λ and nSL selection scans obtained using different datasets. Calculations from nSL analysis are based on the absolute values of the statistic. Genomic region: part of the genome that is included in each dataset. Statistic: selection scan which comparisons were done. Shared outliers: percentage of outliers that are shared between datasets. Shared outliers in inversions: the percentage of outliers shared between datasets that are in inversion regions. Shared outliers in collinear: the percentage of outliers shared between datasets that are in the collinear genome

| Compared datasets | Genomic region | Statistic | Shared outliers | Shared outliers in inversions | Shared outliers in collinear |
| --- | --- | --- | --- | --- | --- |
| SN, N | Inversions + collinear | $\Lambda$ | 23% | 5% | 18% |
|  |  | nSL | 33% | 31% | 2% |
| SNcol, Ncol | Only collinear | $\Lambda$ | 19% | | 19% |
|  |  | nSL | 32% |  | 32% |
| SN, S | Inversions + collinear | $\Lambda$ | 30% | 6% | 24% |
|  |  | nSL | 48% | 39% | 9% |
| SNcol, Scol | Only collinear | $\Lambda$ | 44% | | 44% |
|  |  | nSL | 44% |  | 44% |
| S, N | Inversions + collinear | $\Lambda$ | 9% | 2% | 7% |
|  |  | nSL | 20% | 18% | 2% |
| Scol, Ncol | Only collinear | $\Lambda$ | 24% | | 24% |
|  |  | nSL | 20% |  | 20% |

**Table 4.** Changes in percentage of outliers found in collinear parts of the genome when including or excluding inversions. Results are shown for three datasets (SN, N, and S) analysed using both selection scan methods (Λ and nSL) with results summarised in 50kb windows. Calculations from nSL analysis are based on the absolute values of the statistic. Outliers in collinear regions before excluding inversions: the percentage of outliers in collinear regions identified based on datasets that included inversions, inversions were not removed before identifying the outliers. Lost outliers in collinear parts after excluding inversions: the percentage of outlier windows in collinear parts that disappear after excluding inversions and recalculating the outlier threshold. New outliers in collinear parts after excluding inversions: the percentage of outlier windows in collinear parts that appear after excluding inversions and recalculating the outlier threshold.

| Dataset | Statistic | Outliers in collinear regions before excluding inversions | Lost outliers in collinear parts after excluding inversions | New outliers in collinear parts after excluding inversions |
| --- | --- | --- | --- | --- |
| SN | $\Lambda$ | 83% | 9% | 0% |
|  | nSL | 30% | 0% | 42% |
| S | $\Lambda$ | 48% | 0% | 26% |
|  | nSL | 36% | 0% | 36% |
| N | $\Lambda$ | 60% | 0% | 14% |
|  | nSL | 21% | 0% | 53% |

### Selection scans in inversion homozygotes - detecting false positive and haplotype-specific signals

#### 5. Selection scans in homozygote datasets reveal haplotype-specific and false positive signals

nSL selection signal was detected in ten inversions in both homozygotes, eight inversions in only one homozygote (four of which were minor alleles), and in three inversions selection signals disappeared completely in the homozygous datasets (Table S5). In the Λ scans, three inversions showed selection signals in both homozygotes, and five inversions showed signals in only one homozygote. None of the inversions lost their selection signals in both homozygotes. Figure 6 shows the large variation in the difference between the observed and the expected percentages of outliers among inversions and datasets. For example, Inv2 and Inv16 show a much higher proportion of observed outliers in SNInv and one of the homozygote datasets — dozens observed compared to only a few percent expected. In contrast, Inv13 exhibits similar observed and expected percentages in both SNInv and MJNInv, and notably fewer observed outliers than expected in MJAInv (Figure 6a). Overall, analysis of homozygote datasets revealed that the majority of inversions harbour selection signals in either both or one inversion haplotype. However, in few cases selection signals in homozygote datasets disappeared completely indicating false positive signals obtained in SNInv datasets.

**Figure 6.**
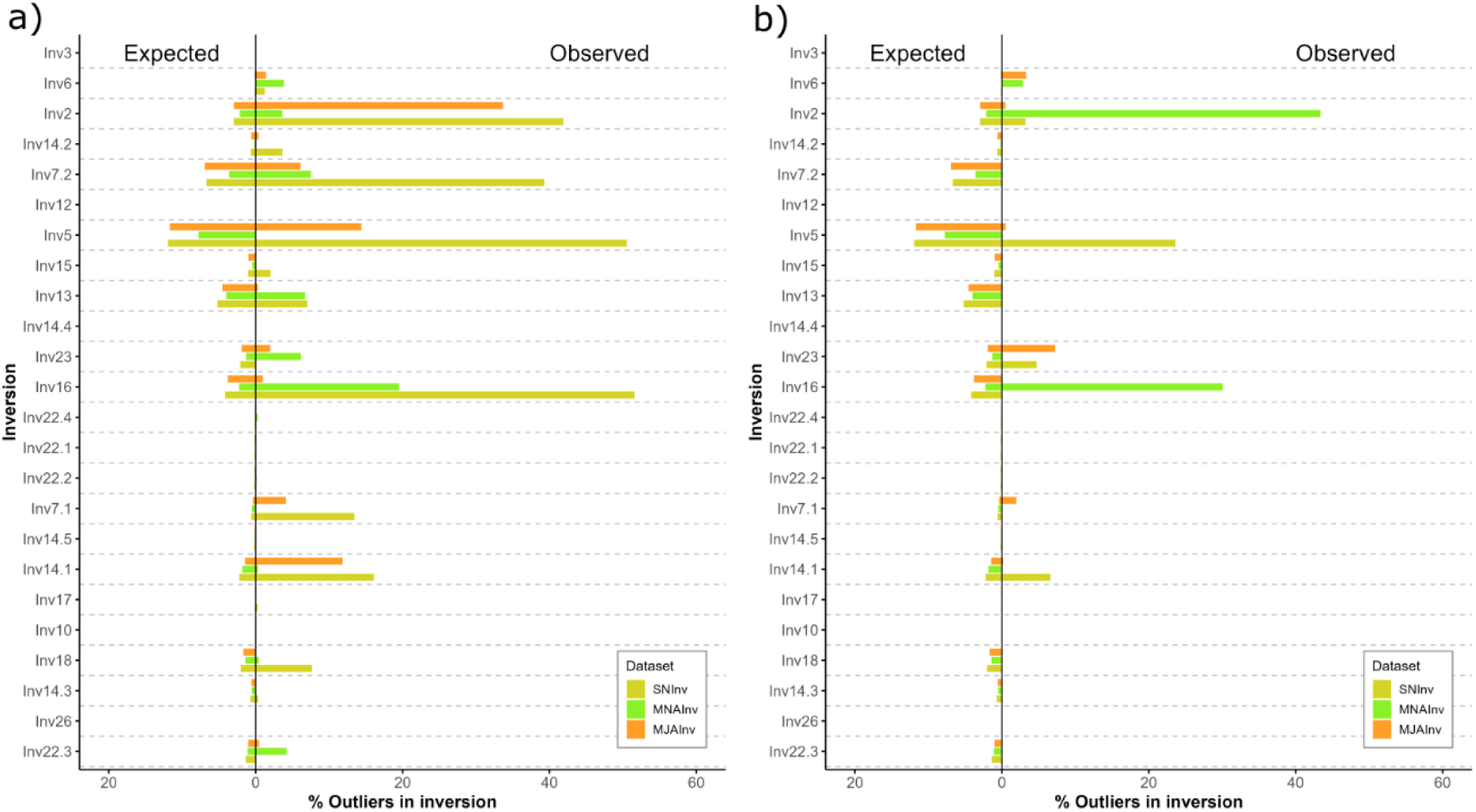
Comparison of the expected and the observed percentages of outliers detected in inversions. The expected values were calculated based on the length of the inversion and a 1% outlier threshold. The results are shown for nSL (a) and Λ (b). All selection scans were performed using 52 SNP windows and three datasets: datasets including individuals carrying any of the possible inversion genotypes (SNInv, yellow); datasets including only individuals that were homozygous for the major allele for the inversion in question (MJAInv, orange); and datasets including only individuals that were homozygous for the minor allele for the inversion in question (MNAInv, green). Each dataset included one analysed inversion and a collinear genome. The inversions are ordered according to the major allele frequency (from lowest to highest, top to bottom).

#### 6. Selection signal tends to be reduced in homozygote datasets

While analyses using SN dataset (all individuals and all inversions), showed that inversion regions are overall enriched for selection signals compared to collinear regions, we observed that the strength of these signals tends to be reduced in homozygote datasets. Our tests indicated a depletion of the signal in homozygotes (MJAInv and MNAInv datasets) compared to the SVInv dataset (Wilcoxon tests, p < 0.02 for both nSL and Λ scans; see Supplementary_Information_Selection_model.html). However, the magnitude of the selection signal in inversion regions did not correlate with inversion haplotype frequency. Similarly, inversion haplotype frequency did not explain the magnitude of selection signals within inversions or indicate any clear patterns regarding what happens to selection signals in inversions (the disappearance of the signal is not dependent on inversion frequency; see Supplementary_Information_Selection_model.html).

In summary, although selection signal tends to be reduced in homozygous datasets compared to SNInv, there is a considerable heterogeneity in this respect among inversions and among datasets (Figure 6, Figure S4, Table S5). The strength of the selection signals and its behaviour in different datasets did not depend on the frequency of inversions.

### Gene ontology terms enriched in selection targets

To explore the functions associated with regions under selection, we performed Gene Ontology (GO) enrichment analysis using outlier windows that (1) were located outside inversions and specific to N population, (2) were located outside inversions and specific to S population, (3) were shared between S, N, and SN datasets, or (4) remained significant after excluding heterozygous individuals in the analysis performed with only homozygotes. In total, we identified 24 significantly enriched GO terms (Table S6). Enriched functions included terms such as “signalling receptor activity” and “response to stimulus,” potentially related to odorant receptor function. A complete list of enriched GO terms, including fold enrichment and significance levels, is provided in Table S6.

## Discussion

In this study we evaluate performance of two selection scan methods using simulated datasets and used whole-genome resequencing data from 312 individuals of the spruce bark beetle to investigate how selection signals are distributed throughout its genome and how chromosomal inversions influence these patterns and our ability to detect them.

Selection scan results based on the simulated datasets indicated that the two evaluated statistics differed in their sensitivity to suppressed recombination in the inversion regions. As expected, Λ was highly sensitive to low recombination [12] and produced distinct outliers in the majority of simulation-based selection scans (Figure 3 and Supplementary Materials: Simulation Results .zip folder). In contrast, nSL, consistent with its known robustness to recombination variation [27], yielded much less pronounced outlier peaks in inversions, though a low level of background noise remained, with some outliers detected even in neutrally evolving regions. Importantly, these nSL outliers did not form visible clusters and usually appeared as single-locus signals. Additionally, simulations showed that analysing homozygote-only datasets can substantially reduce false-positive signals indicating that this approach can be valuable in study systems where inversion genotyping is possible.

As predicted, we observed a significant enrichment of selection signals within inversion regions using both nSL and Λ statistics in the spruce bark beetle data. Notably, even in the homozygotes, most inversions exhibit some selection signals in at least one homozygote dataset. These findings reinforce the view that most inversions contribute to adaptation [42,68–69] and are unlikely to be selectively neutral [31], particularly in the case of large polymorphic inversions [36] — the kind identified in the spruce bark beetle genome [50].

On the other hand, we observed substantial variation in both the magnitude and behaviour (enrichment or depletion) of selection signals across inversions and datasets, in particular when all genotypes were contrasted with homozygotes. In general, in the concordance with simulation-based results, selection signals were weaker in the homozygote datasets, suggesting that some signals detected in the entire dataset may be significantly inflated by the presence of heterozygous individuals. By restricting analyses to individuals carrying only one inversion haplotype, we reduced the likelihood of false positives and were able to detect potential haplotype-specific selection signals. However, the small sample size of some homozygote datasets (directly reflecting inversion frequency) could limit the ability to detect selection signals. Our simulation-based tests indicated that small sample sizes may lead to underestimation of outliers detected in inversions. Overall, the lack of a clear relationship between the extent of inversion polymorphism and the behaviour of selection signals across the spruce bark beetle datasets underscores a key challenge: although inversions are enriched for selection signals and homozygote datasets can help in reducing false positive signals, identifying the specific targets of selection within these regions remains difficult.

Contrary to our expectations, we found that the presence of inversions had only a minimal impact on the shape of the selection landscape and our ability of detect selection signals in the collinear genome. Importantly, this funding suggests that identifying loci under selection outside of inversions remains feasible, even in genomes characterized by extensive inversion polymorphism. This result aligns with our earlier findings on demographic inference [56], where we showed that overlooking chromosomal inversions had no effect on model selection and introduced only minor biases in the estimation of demographic parameters.

Additionally, the spruce bark beetle selection scans supported the necessity of using multiple complementary approaches to selection scans. While the overall patterns of selection inferred by Λ and nSL were similar, differences were observed at a local scale. There was a high correlation of selection signals at 50 kb windows, but only a limited overlap among outlier windows at 52 SNPs, which demonstrates the general difficulty of selection scans in pinpointing the focal gene under selection [70]. Moreover, the selection of window sizes in the Λ scans can influence which regions are considered candidates for selection. The differences observed in the spruce bark beetle between 52 and 117 SNP windows can, to some extent, reflect differences in the ability to detect more recent and more ancient sweep signatures: smaller window sizes are more effective at detecting ancient sweeps [12].

Our study revealed a broadly similar selection landscape between the southern and northern populations of the spruce bark beetle, suggesting that these populations are likely subject to comparable selective pressures and/or that their divergence is relatively recent, with the observed signals possibly reflecting selection in a shared ancestral population. According to our recent demographic modelling, the divergence between the southern and northern clades occurred approximately 80,000 years ago [56]. Signals of selection in the site frequency spectrum are expected to persist for roughly 0.1Ne generations, while haplotype-based signals fade more rapidly, typically within ∼0.01Ne generations [71–72]. Assuming an ancestral Ne of 250,000 [56], haplotype-based signatures could persist for around 2,500 generations—well after the estimated divergence time—supporting the idea that shared selection pressures may be acting across both populations. The inferred Ne for the northern and southern populations after the split was 180,000 and 830,000, respectively, and even under the higher Ne scenario, the timescale of detectable selection signals remains post-divergence. These findings suggest that spruce bark beetle populations across Europe may be subject to similar selective pressure. Nonetheless, we also detected evidence of regional differentiation: one of the strongest selection signals outside of inversions (on IpsContig3) overlapped with a pronounced Fst peak between the northern and southern groups, indicating that divergent selection may also contribute to shaping the selection landscape in this species.

Finally, our GO enrichment analysis identified terms such as “signalling receptor activity” and “response to stimulus,” which are potentially linked to odorant receptor function. These findings are of particular interest considering recent studies showing that inversions are enriched in genes encoding odorant receptors [50], and that these inversions may contain receptor variants that respond differently to different components of pheromones [73].

## Conclusions and future directions

Our study provides new insights into the selection landscape across the spruce bark beetle genome, identifying candidate genes that may guide future research into the genomic basis of adaptation in this ecologically and economically important species. We show that most polymorphic inversions in the genome are enriched for selection signals, supporting their role as key contributors to adaptation, while also highlighting the importance of accounting for structural variation — particularly inversions — when performing selection inference. As illustrated by the changing selection patterns in the homozygote datasets in simulations as well as empirical datasets, inversions can affect the detection of selection within their regions. Nevertheless, their presence does not obscure the identification of selection signals outside inversions, emphasizing the feasibility of conducting genome-wide selection scans even in inversion-rich genomes. More broadly, our results underscore the need to systematically evaluate the performance and robustness of evolutionary inference methods in the presence of structural variation [15,28]. However, such efforts remain rare, and further research is needed to understand how inversions influence evolutionary inference, particularly given that their effects likely vary across species with differing numbers and evolutionary histories of large polymorphic inversions.

## Acknowledgments

We thank members of the Genomics and Experimental Evolution Group at Jagiellonian University for their help in improving this manuscript. This work was funded by a Polish National Science Center 2018/30/E/NZ8/00105 grant to K.N.B. We gratefully acknowledge Polish high-performance computing infrastructure PLGrid (HPC Centers: ACK Cyfronet AGH) for providing computer facilities and support within computational grant no. PLG/2024/017278 to K.N.B.

## Data Availability

All DNA sequences have been deposited in the National Centre for Biotechnology Information Sequence Read Archive under BioProject ID PRJNA1013983. Scripts used for data processing, generation of intermediate files, and analyses, have been archived and are publicly available in the Dryad Digital Repository entry [will be added as soon as possible].

## Author contributions

J.M.G., K.N.B. conceived the study and wrote the first draft of the manuscript. J.M.G, performed all the main analyses. P.Z processed and supervised the raw data processing. W.B. performed the GO analysis. W.B. and P.Z helped in data interpretation and provided feedback on all manuscript versions. All authors read and approved the final manuscript.

