## Supplementary Information for "Selection inference in a complex genomic landscape: the impact of polymorphic inversions"

1. Correlations and permutations

In this section, we present correlation analysis conducted in the study. For each analysis, we computed the observed correlation coefficient (r, Pearson's r when both variables are binary, and Spearman's otherwise) and report its squared value ($r^2$, in Tables). To assess significance of the correlations, we performed 1000 permutations (permuting one of the variables that we correlate). All corresponding density plots display the distribution of correlation coefficient obtained using permuted datasets with dashed lines indicating the value of correlation coefficient obtained for the original data (Obs r). The mean of the correlation coefficient obtained from permuted datasets is also presented at the plots (Mean perm r).

1.1 Correlation between genomic region (inversion vs. collinear) and selection outlier status (outlier vs. non-outlier)

To assess whether selection outliers tend to occur more frequently in inverted than in collinear genomic regions, we computed Pearson's correlations between two binary variables at the window level: (i) genomic region, encoded as 0 for collinear and 1 for inverted windows; and (ii) outlier status, encoded as 0 for non-outlier windows and 1 for outlier windows. This analysis was performed based on both nSL and Λ selection scan results, using two SNP window sizes (52 and 117 SNPs). Observed correlations were significantly higher than expected ($p < 0.001$ in all cases except one), supporting an enrichment of outlier SNPs in inversion regions and suggesting that selection signals are non-randomly distributed across the genome.

Table S1.1. Pearson correlation coefficients (r), ($r^2$), and p-values for the association between genomic region (collinear vs. inverted) and selection outlier status (non-outlier vs. outlier) across three datasets (SN, S, N), for Λ and nSL statistics, using 52 and 117 SNP window sizes. Asterisks indicate statistically significant results ($p < 0.001$).

| Dataset | Window size | Statistic | r | $r^2$ | p-value |
|---|---|---|---|---|---|
| SN | 52 | Λ | -0.0055 | 0 | >0.05 |
| SN | 52 | nSL | 0.1161 | 0.0135 | <0.001* |
| SN | 117 | Λ | 0.1490 | 0.0222 | <0.001* |
| SN | 117 | nSL | 0.1145 | 0.0131 | <0.001* |
| S | 52 | Λ | 0.0848 | 0.0072 | <0.001* |
| S | 52 | nSL | 0.1201 | 0.0144 | <0.001* |
| S | 117 | Λ | 0.1491 | 0.0222 | <0.001* |
| S | 117 | nSL | 0.1205 | 0.0145 | <0.001* |
| N | 52 | Λ | 0.0519 | 0.0027 | <0.001* |
| N | 52 | nSL | 0.1265 | 0.0160 | <0.001* |
| N | 117 | Λ | 0.1468 | 0.0216 | <0.001* |
| N | 117 | nSL | 0.1320 | 0.0174 | <0.001* |

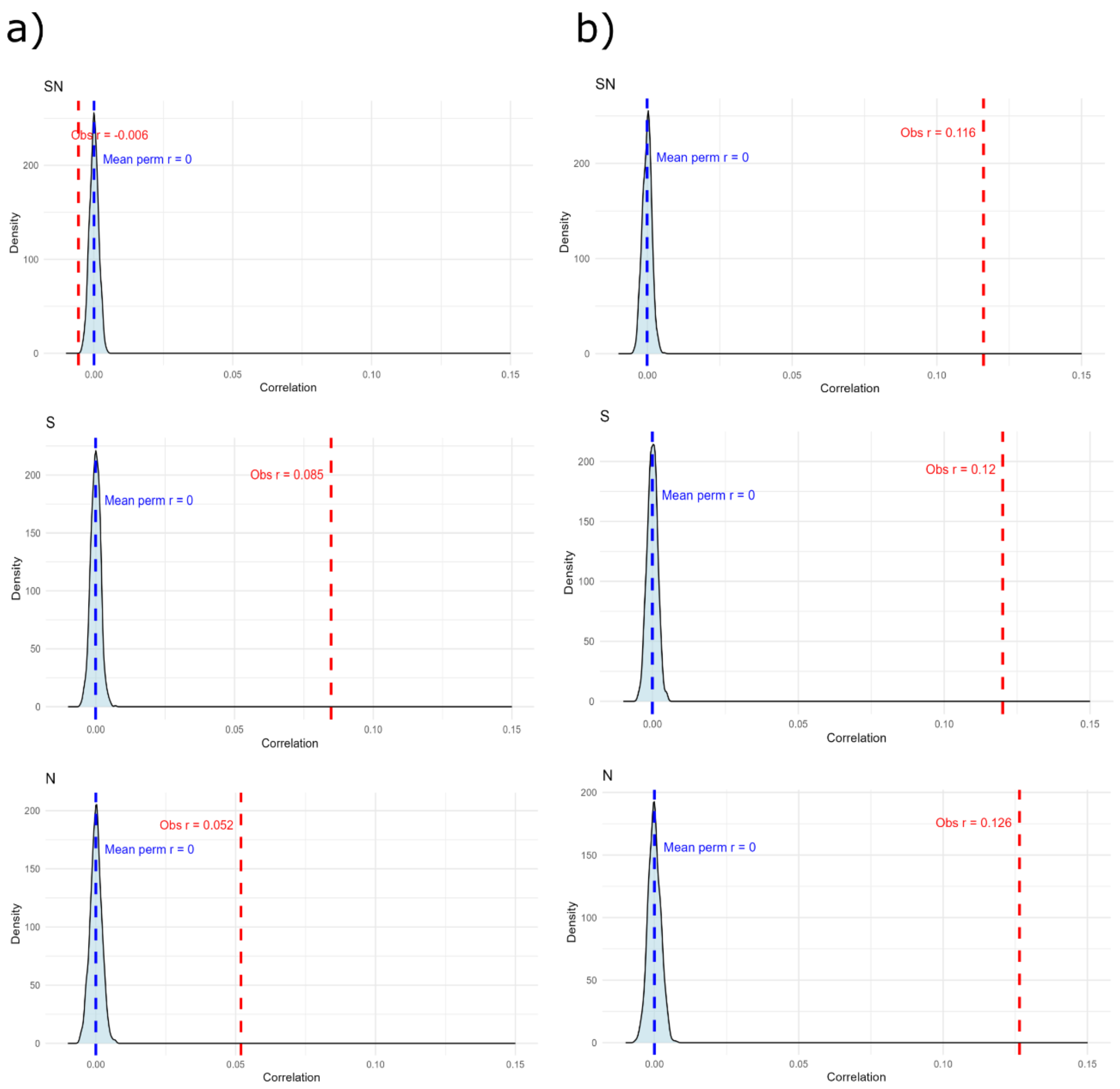


Figure S1.1.I Permutation density plots of correlation coefficients for Λ (a) and nSL (b) selection scans with 52 SNPs window size across three population datasets SN (top row), S (middle row), and N (bottom row). In each panel, the light blue shaded area represents the null distribution of Pearson's correlation coefficients calculated from 1000 permutations of the data; the blue dashed line marks the mean correlation coefficient for the permuted datasets ("Mean perm r"), and the red dashed line indicates the observed correlation ("Obs r").

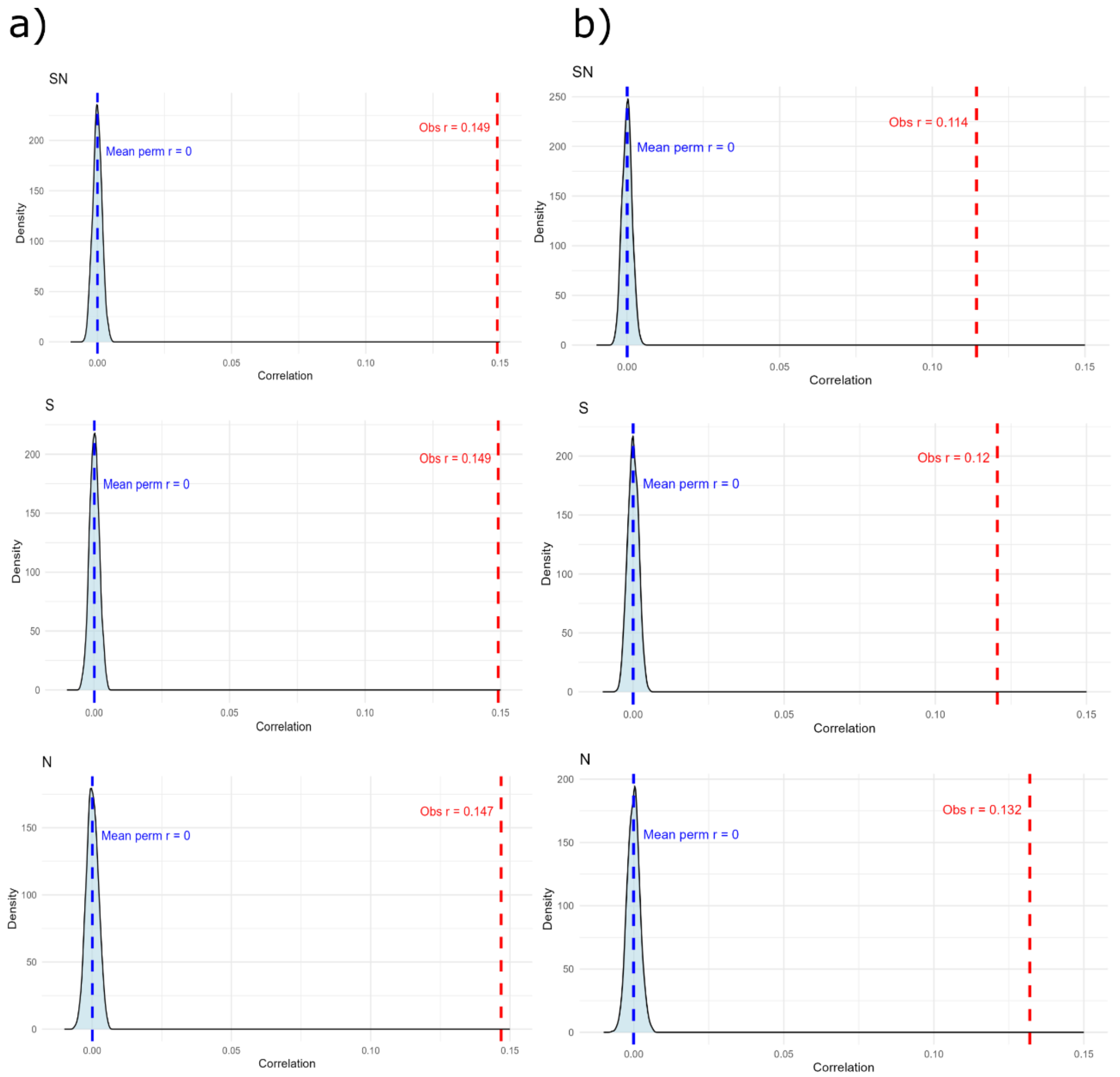


Figure S1.1.II Permutation density plots of correlation coefficients for Λ (a) and nSL (b) selection scans with 117 SNPs window size across three population datasets SN (top row), S (middle row), and N (bottom row). In each panel, the light blue shaded area represents the null distribution of Pearson's correlation coefficients calculated from 1000 permutations of the data; the blue dashed line marks the mean correlation coefficient for the permuted datasets ("Mean perm r"), and the red dashed line indicates the observed correlation ("Obs r"). Higher observed values (red) located to the right of the null peak (blue) show stronger than expected associations under the permutation test.

#### 1.2 Correlation between Fst values and selection scan results

To test whether genetic differentiation is associated with the strength of the selection signals, we computed Spearman correlations between Fst values calculated between southern and northern populations and selection scans results obtained in both nSL and Λ selection scans. This analysis was performed using both 52 and 117 SNP window sizes. We generated a null distribution of correlation coefficients by permuting the selection scans results 1000 times while keeping Fst values fixed, under the null hypothesis of no association between Fst values and the magnitude of the selection signals. The observed correlation was weak but significantly higher than expected by chance ($p < 0.001$), indicating that windows with higher genetic differentiation (Fst) also tend to exhibit stronger selection signals.

Table S1.2. Spearman correlation coefficients (r), ($r^2$), and p-values for the association between genetic differentiation (Fst) and selection scan results (Λ and nSL) across three datasets (SN, S, N), using 52 and 117 SNP window sizes. Asterisks indicate statistically significant results ($p < 0.001$).

| Dataset | Window size | Statistic | r | $r^2$ | p-value |
|---|---|---|---|---|---|
| SN | 52 | Λ | 0.0205 | 0.0004 | <0.001* |
| SN | 52 | nSL | 0.0683 | 0.0047 | <0.001* |
| SN | 117 | Λ | 0.1694 | 0.0287 | <0.001* |
| SN | 117 | nSL | 0.1069 | 0.0114 | <0.001* |
| S | 52 | Λ | 0.0406 | 0.0016 | <0.001* |
| S | 52 | nSL | 0.0783 | 0.0061 | <0.001* |
| S | 117 | Λ | 0.1442 | 0.0208 | <0.001* |
| S | 117 | nSL | 0.1136 | 0.0129 | <0.001* |
| N | 52 | Λ | 0.1989 | 0.0396 | <0.001* |
| N | 52 | nSL | 0.1256 | 0.0158 | <0.001* |
| N | 117 | Λ | 0.2518 | 0.0634 | <0.001* |
| N | 117 | nSL | 0.1614 | 0.0260 | <0.001* |

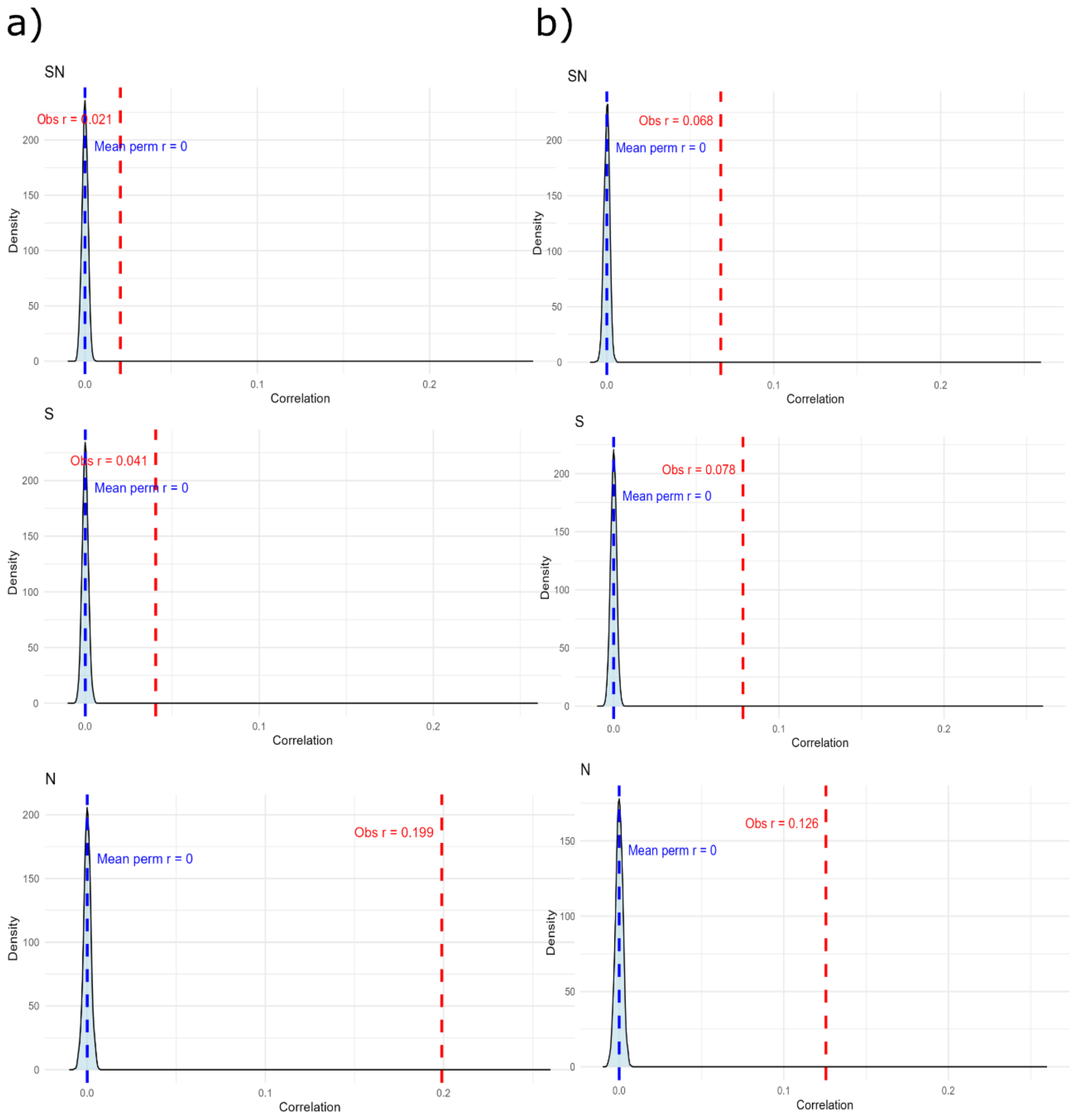


Figure S1.2.I Permutation density plots of Spearman correlation coefficients between Fst and selection scans computed using 52 SNP window size in three population datasets: SN (top row), S (middle row), and N (bottom row). Panels (a) show results for Λ; panels (b) for nSL. In each panel, the light blue shaded area represents the null distribution of Spearman's correlation coefficients calculated from 1000 permutations of the data; the blue dashed line marks the mean correlation coefficient for the permuted datasets ("Mean perm r"), and the red dashed line indicates the observed correlation ("Obs r"). In all cases, observed correlations were significant.

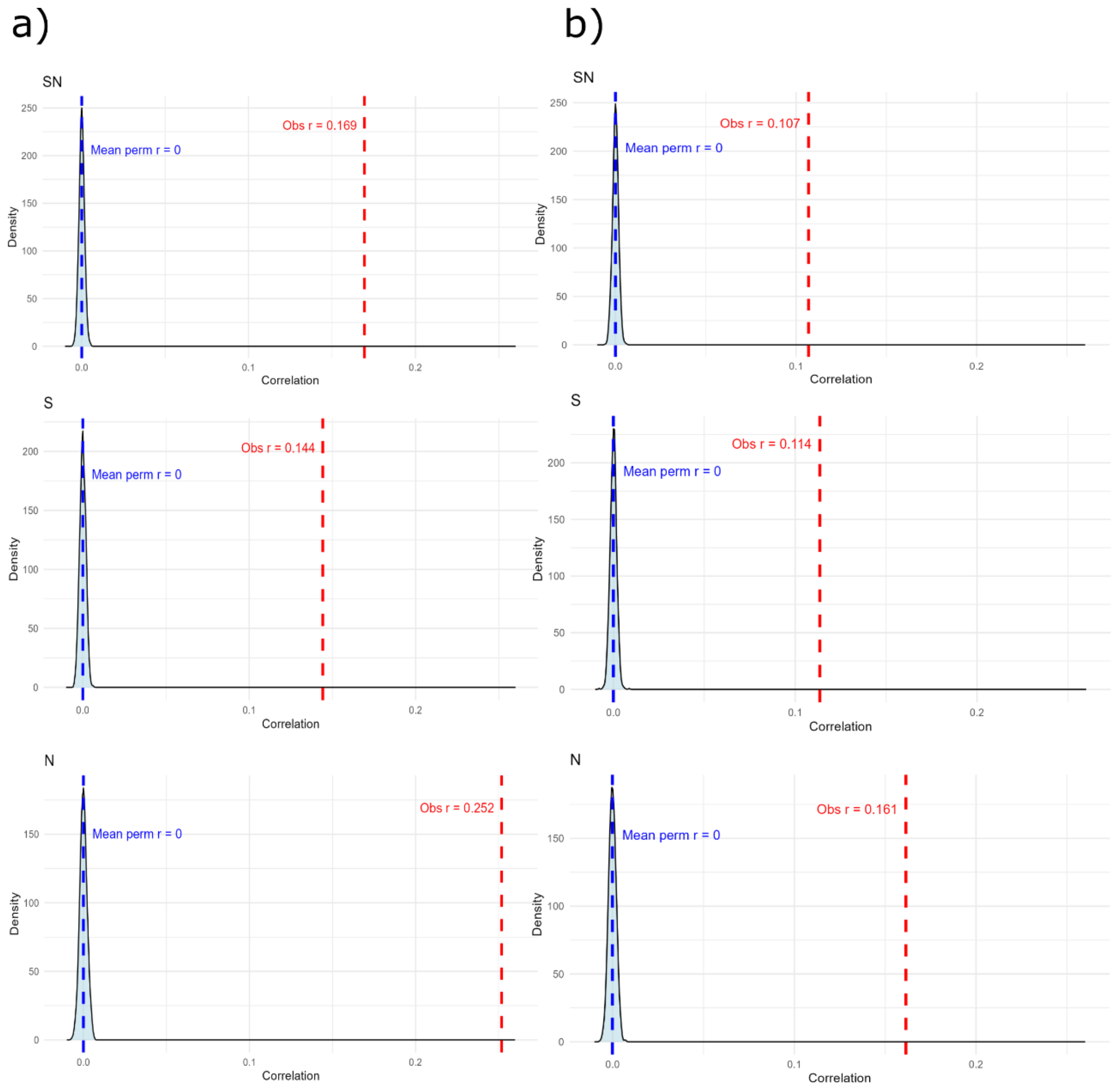


Figure S1.2.II Permutation density plots of Spearman correlation coefficients between Fst and selection scans computed using 117 SNP window size in three population datasets: SN (top row), S (middle row), and N (bottom row). Panels (a) show results for Λ; panels (b) for nSL. In each panel, the light blue shaded area represents the null distribution of Spearman's correlation coefficients calculated from 1000 permutations of the data; the blue dashed line marks the mean correlation coefficient for the permuted datasets ("Mean perm r"), and the red dashed line indicates the observed correlation ("Obs r").

#### 1.3 Correlation between inversion haplotype frequency and selection scan results

To test whether the frequency of inversion haplotypes correlates with the strength of selection signals, we computed Spearman correlations between haplotype frequency and selection scan results. Specifically, we calculated the correlation between inversion haplotype frequencies (major: MJA; minor: MNA) and the mean value of the selection scan statistic within each inversion obtained for the SN dataset. The hypothesis is that inversions with intermediate frequencies where heterozygotes are most frequent should exhibit stronger selection signals due to recombination suppression. Null distributions of correlations coefficients were obtained by permutating frequency 1000 times and calculating correlation coefficient for each of the permuted dataset. The observed correlations were not significant, so we found no correlation between the inversion haplotype frequency and the magnitude of the selection scans results.

Table S1.3. Spearman correlation coefficients (r), ($r^2$), and p-values for the association between inversion haplotype frequency (major (MJA) and minor (MNA)) and selection scan results (Λ and nSL) in the SN dataset, using 52 and 117 SNP window sizes. Asterisks indicate statistically significant results ($p < 0.001$).

| Dataset | Window size | Statistic | r MJA | $r^2$ MJA | r MNA | $r^2$ MNA | p-value |
|---|---|---|---|---|---|---|---|
| SN | 52 | Λ | 0.0700 | 0.0049 | -0.0465 | 0.0022 | >0.05 |
| SN | 52 | nSL | -0.0639 | 0.0041 | 0.1057 | 0.0112 | >0.05 |
| SN | 117 | Λ | 0.0966 | 0.0093 | -0.0635 | 0.0040 | >0.05 |
| SN | 117 | nSL | -0.0887 | 0.0079 | 0.1348 | 0.0182 | >0.05 |

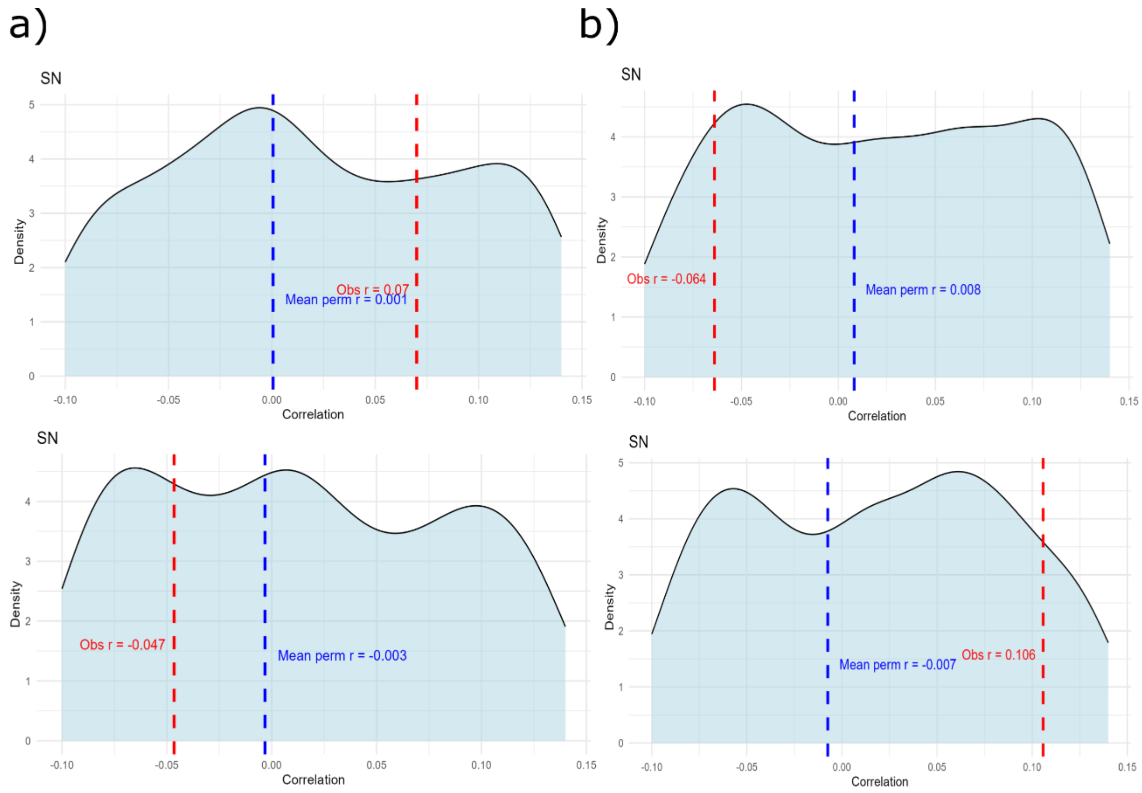


Figure S1.3.I Permutation density and scatter plots of Spearman correlation coefficients between inversion haplotype frequency and selection scans results using 52 SNPs window size. Both rows show permutation distributions of Spearman correlation coefficients between a) Λ and b) nSL values and the frequency of the major haplotype (MJA, top row) or the minor haplotype (MNA, bottom row). Light blue areas represent the null distribution obtained from 1000 permutations. Red dashed lines indicate the observed correlation ("Obs r") and blue dashed lines the mean correlation coefficient calculated based on the permutations ("Mean perm r").

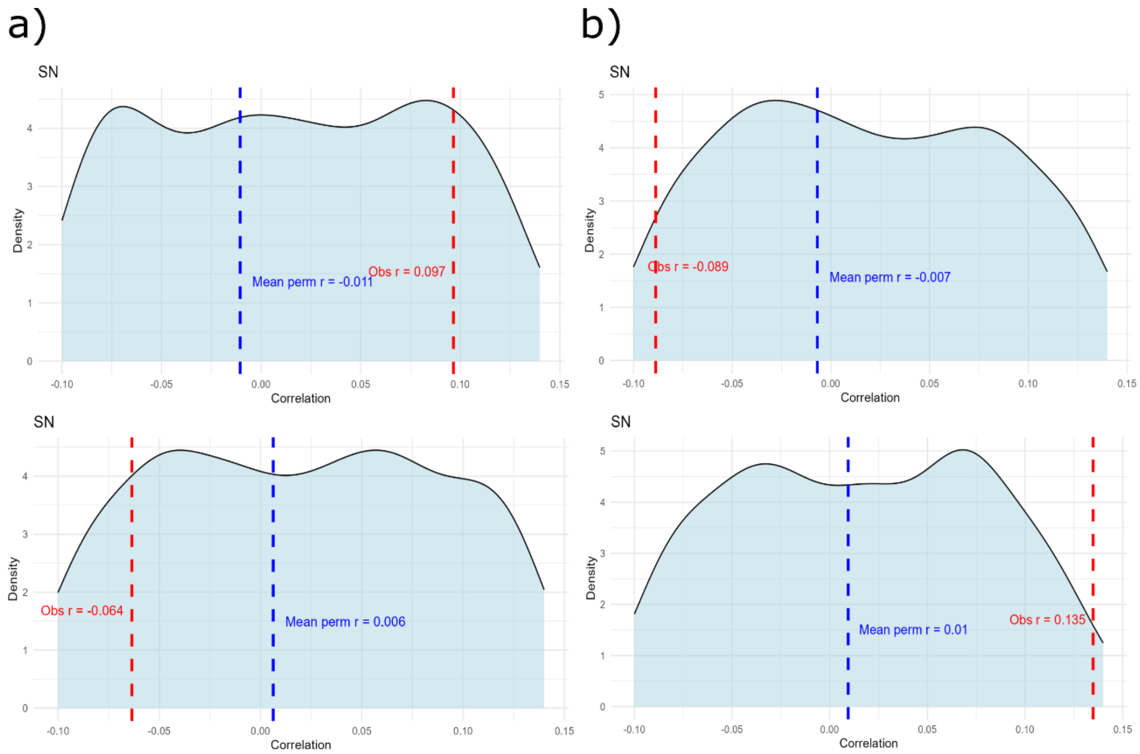


Figure S1.3.II Permutation density and scatter plots of Spearman correlation coefficients between inversion haplotype frequency and selection scans results using 117 SNPs window size. Both rows show permutation distributions of Spearman correlation coefficients between a) Λ and b) nSL values and the frequency of the major haplotype (MJA, top row) or the minor haplotype (MNA, bottom row). Light blue areas represent the null distribution obtained from 1000 permutations. Red dashed lines indicate the observed correlation ("Obs. r") and blue dashed lines the mean correlation coefficient calculated based on the permutations ("Mean perm r").

### 1.4 Correlation between nSL and Λ selection scans results

To assess the consistency between different selection scan approaches, we computed Spearman correlations between nSL and Λ values. This analysis was performed using 52 and 117 SNP window size. We generated a null distribution of correlation coefficients obtained by permuting the nSL values 1000 times while keeping Λ fixed and calculating correlation coefficient for each of the permuted dataset. The observed correlation coefficient was moderate and significant ($p < 0.001$), indicating that the obtained selection landscapes in both nSL and Λ selection scans were similar.

Table S1.4. Spearman correlation coefficients (r), ($r^2$), and p-values for the association between Λ and nSL selection scan statistics across three datasets (SN, S, N), using 52 and 117 SNP window sizes. Asterisks indicate statistically significant results ($p < 0.001$).

| Dataset | Window size | r | $r^2$ | p-value |
|---|---|---|---|---|
| SN | 52 | 0.2969 | 0.0882 | <0.001* |
| SN | 117 | 0.4417 | 0.1951 | <0.001* |
| S | 52 | 0.3515 | 0.1235 | <0.001* |
| S | 117 | 0.4533 | 0.2055 | <0.001* |
| N | 52 | 0.4333 | 0.1878 | <0.001* |
| N | 117 | 0.4099 | 0.1681 | <0.001* |

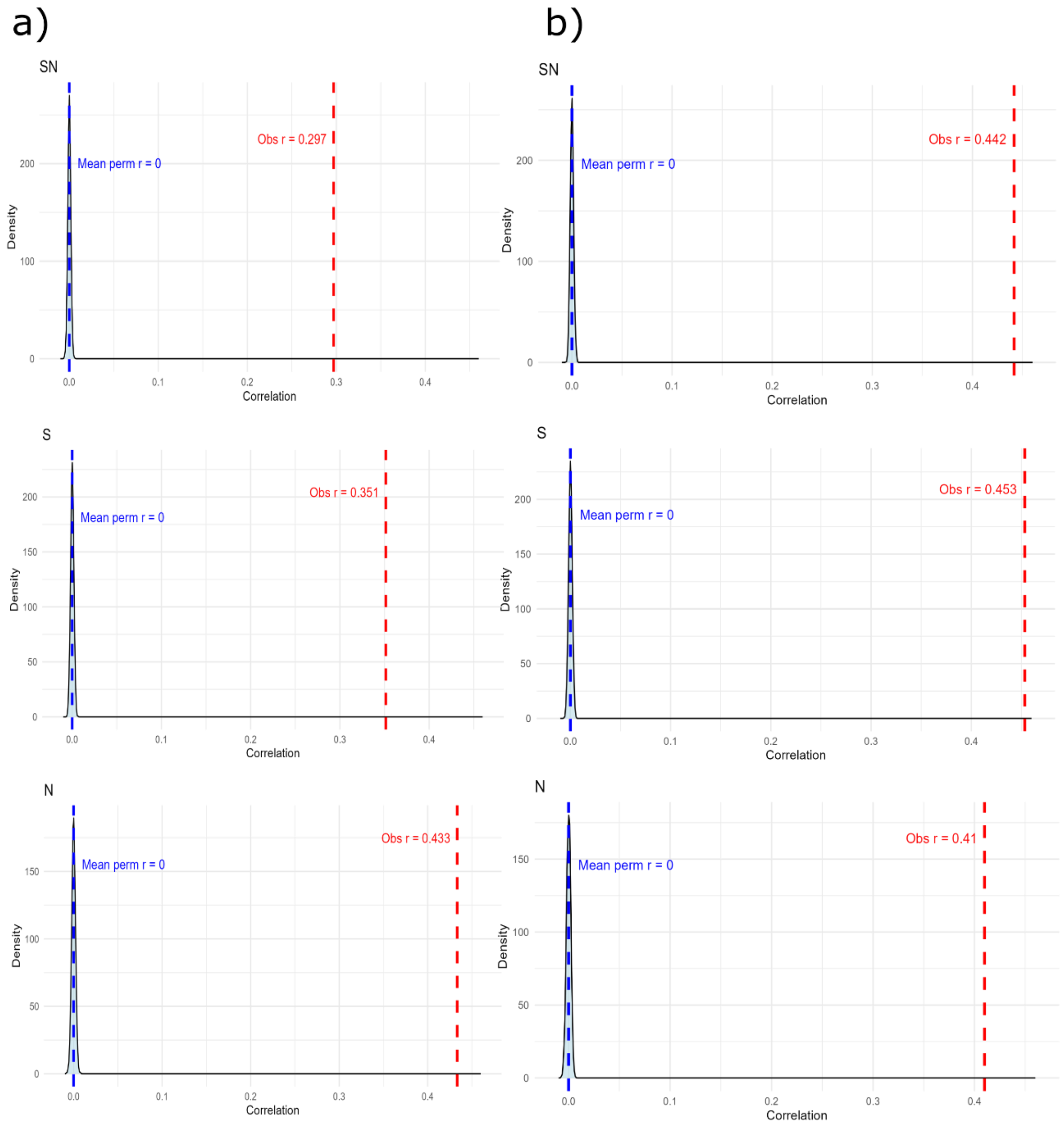


Figure S1.4 Permutation density plots of Spearman correlation coefficients between nSL and Λ selection scans results computed using (a) 52 and (b) 117 SNP window size in three population datasets: SN (top row), S (middle row), and N (bottom row). In each panel, the light blue shaded area represents the null distribution of Spearman's correlation coefficients calculated from 1000 permutations of the data; the blue dashed line marks the mean correlation coefficient obtained from the permuted datasets ("Mean perm r"), and the red dashed line indicates the observed correlation ("Obs r").

### 1.5 Correlation between S and N populations selection signals

To compare the distribution of selection signals across S and N populations, we computed Spearman correlations between Λ and nSL values from each population. Because the number of windows differs between datasets, we used selection scans results summarised in 50kb windows for both 52 and 117 SNPs window sizes. We generated a null distribution of correlation coefficients obtained by permuting 1000 time the results for the S population. The observed correlation coefficient was high and significant, suggesting that the same genomic regions are under selection in both populations.

Table S1.5. Spearman correlation coefficients (r), ($r^2$), and p-values for the association between selection scan values (Λ and nSL) obtained in S and N populations, using 52 and 117 SNP window sizes. Asterisks indicate statistically significant results ($p < 0.001$).

| Selection scan | Window size | r | $r^2$ | p-value |
|---|---|---|---|---|
| Λ | 52 | 0.7645 | 0.5845 | <0.001* |
| nSL | 52 | 0.8075 | 0.6520 | <0.001* |
| Λ | 117 | 0.8097 | 0.6557 | <0.001* |
| nSL | 117 | 0.7945 | 0.6312 | <0.001* |

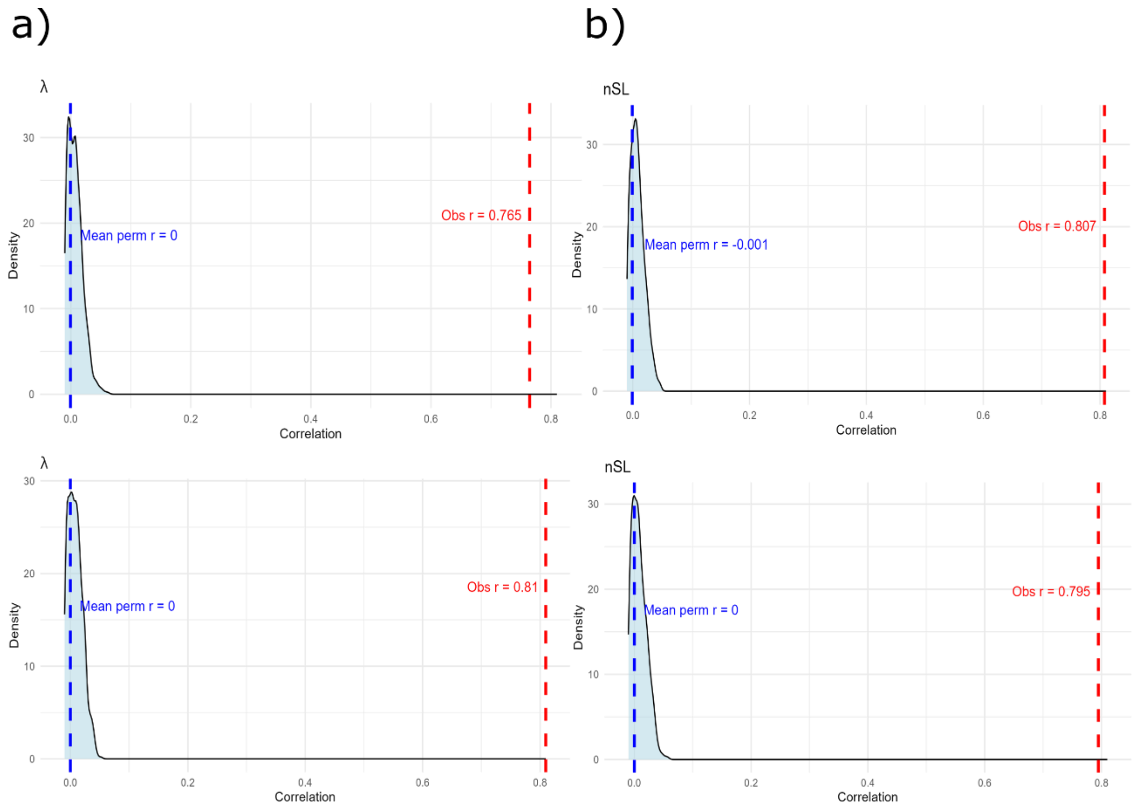


Figure S1.5 Permutation density plots of Spearman correlation coefficients between S and N populations in both selection scans (a) Λ and (b) nSL and 52 (top row) and 117 (bottom row) SNP window size. In each panel, the light blue shaded area represents the null distribution of Spearman's correlation coefficients calculated from 1000 permutations of the data; the blue dashed line marks the mean correlation coefficient obtained from the permuted datasets ("Mean perm r"), and the red dashed line indicates the observed correlation ("Obs r").

#### 1.6 Correlation between selection results that included inverted and collinear regions and results that included only collinear regions

To evaluate the impact of inversion regions on the inference of selection in collinear parts of the genome, we computed Spearman correlations between selection scans results from datasets that included both inverted and collinear regions with those that only included collinear regions. We did it for both selection scans, Λ and nSL, and SNP window sizes (52 and 117) using selection scans results summarised in 50kb windows. We generated a null distribution of correlation coefficients obtained by permuting 1000 times the results obtained using the dataset that included both inverted and collinear parts. The observed correlation indicates that both datasets are highly correlated, suggesting that the presence of inversions has minimal impact on the detection of selection signals in collinear regions.

Table S1.6. Spearman correlation coefficients (r), ($r^2$), and p-values for the association between selection scan results obtained in whole-genome datasets (inversions + collinear regions) and datasets including only collinear regions, across three datasets (SN, S, N), for Λ and nSL statistics, using 52 and 117 SNP window sizes. Asterisks indicate statistically significant results ($p < 0.001$).

| Dataset | Window size | Statistic | r | $r^2$ | p-value |
|---|---|---|---|---|---|
| SN | 52 | Λ | 0.9723 | 0.9454 | <0.001* |
| SN | 52 | nSL | 0.9512 | 0.9047 | <0.001* |
| SN | 117 | Λ | 0.8709 | 0.7584 | <0.001* |
| SN | 117 | nSL | 0.9396 | 0.8829 | <0.001* |
| S | 52 | Λ | 0.9331 | 0.8706 | <0.001* |
| S | 52 | nSL | 0.9157 | 0.8385 | <0.001* |
| S | 117 | Λ | 0.8913 | 0.7945 | <0.001* |
| S | 117 | nSL | 0.8908 | 0.7934 | <0.001* |
| N | 52 | Λ | 0.8916 | 0.7950 | <0.001* |
| N | 52 | nSL | 0.9688 | 0.9386 | <0.001* |
| N | 117 | Λ | 0.9238 | 0.8535 | <0.001* |
| N | 117 | nSL | 0.9673 | 0.9357 | <0.001* |

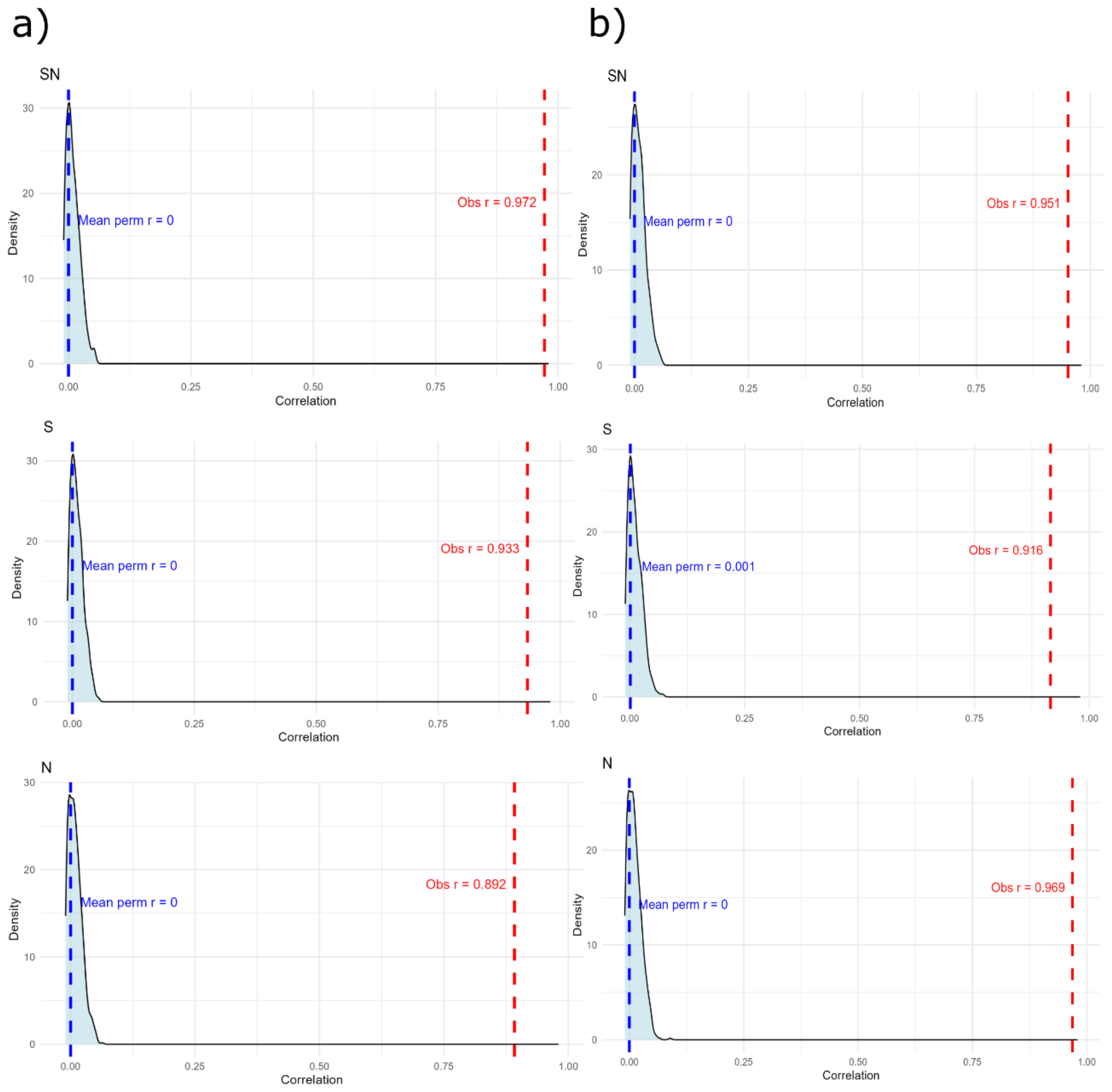


Figure S1.6.I Permutation density plots of Spearman correlation coefficients between selections scans results from datasets that included inverted and collinear regions and datasets that included only collinear regions of the genome. Those results are from selection scans that were performed with 52 SNP window size. The plots show results from selection scans using three population datasets: SN (top row), S (middle row), and N (bottom row). Panels (a) show results for Λ; panels (b) for nSL. In each panel, the light blue shaded area represents the null distribution of Spearman's correlation coefficients calculated from 1000 permutations of the data; the blue dashed line marks the mean correlation from the permuted datasets ("Mean perm r"), and the red dashed line indicates the observed correlation ("Obs r").

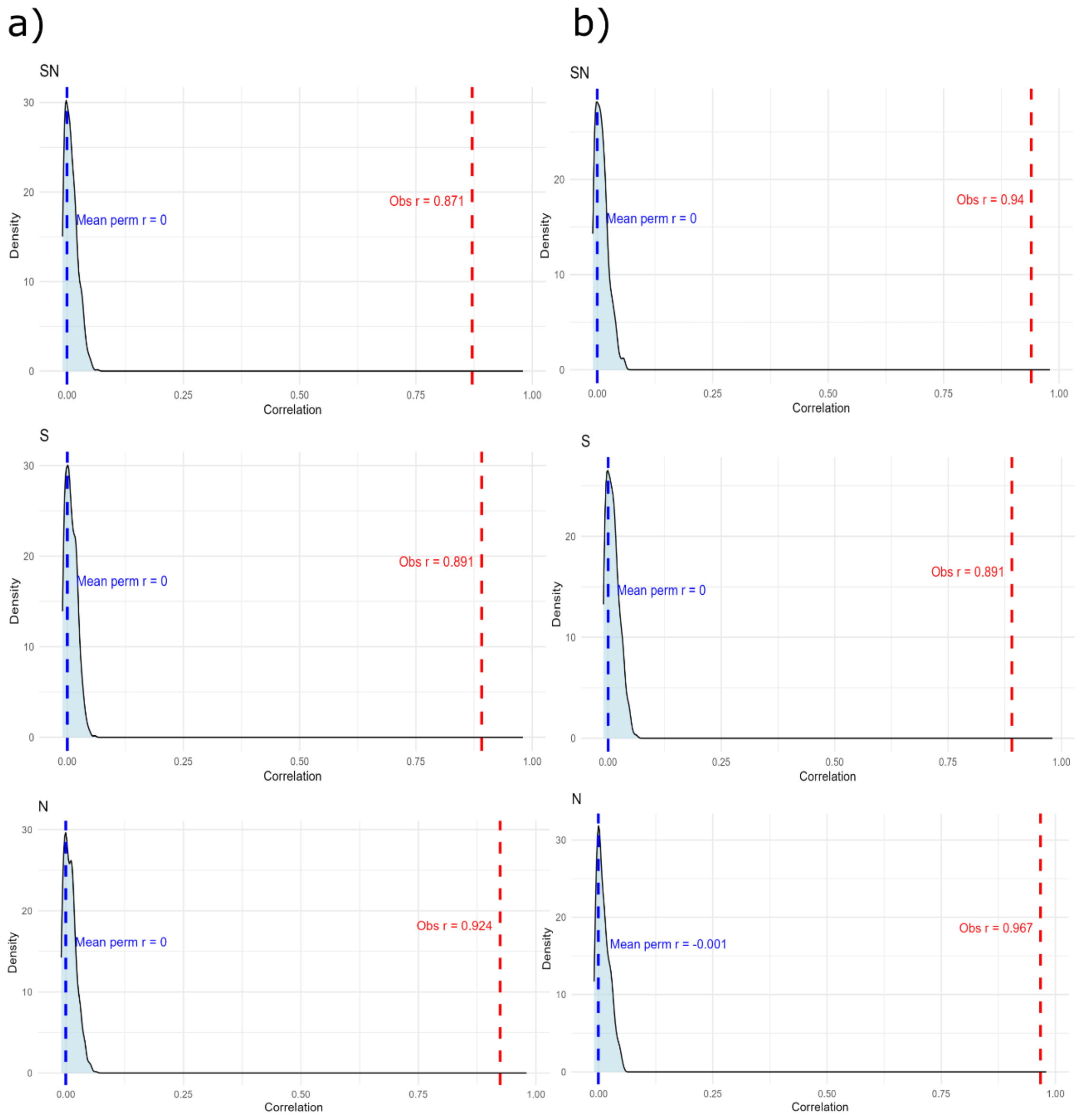


Figure S1.6.II Permutation density plots of Spearman correlation coefficients between selections scans results from datasets that included inverted and collinear regions and datasets that included only collinear regions of the genome. Those results are from selection scans that were performed with 117 SNP window size. The plots show results from selection scans using three population datasets: SN (top row), S (middle row), and N (bottom row). Panels (a) show results for Λ; panels (b) for nSL. In each panel, the light blue shaded area represents the null distribution of Spearman's correlation coefficients calculated from 1000 permutations of the data; the blue dashed line marks the mean correlation from the permuted datasets ("Mean perm r"), and the red dashed line indicates the observed correlation ("Obs r").

### 1.7 Correlation between the status of the windows (outlier or non-outlier) in selection scans results obtained in two selection scan methods

To assess whether different selection scan methods identify the same genomic regions under selection, we computed Pearson correlations between the binary outlier classifications from nSL and Λ selection scans. Each genomic window was assigned a value of 1 (outlier) or 0 (non-outlier) for each method. We generated a null distribution of correlation coefficients obtained by permuting the column with outlier information for nSL 1000 times. The correlations were weak but significantly higher than expected. Only for the southern population with 117 SNP windows, the correlation between nSL and Λ outlier window status was not significant ($p > 0.05$, based on 1000 data permutations). Despite using different methods to detect selection, both nSL and Λ scans tend to identify similar genomic regions as outliers.

Table S1.7. Pearson correlation coefficients (r), ($r^2$), and p-values for the association between outlier classifications from Λ and nSL selection scans across three datasets (SN, S, N), using 52 and 117 SNP window sizes. Asterisks indicate statistically significant results ($p < 0.001$).

| Dataset | Window size | r | $r^2$ | p-value |
|---|---|---|---|---|
| SN | 52 | 0.1505 | 0.0227 | <0.001* |
| S | 52 | 0.1906 | 0.0363 | <0.001* |
| N | 52 | 0.1620 | 0.0262 | <0.001* |
| SN | 117 | 0.0094 | 1e-04 | <0.001* |
| S | 117 | -0.0033 | 1e-05 | >0.05 |
| N | 117 | 0.2725 | 0.0743 | <0.001* |

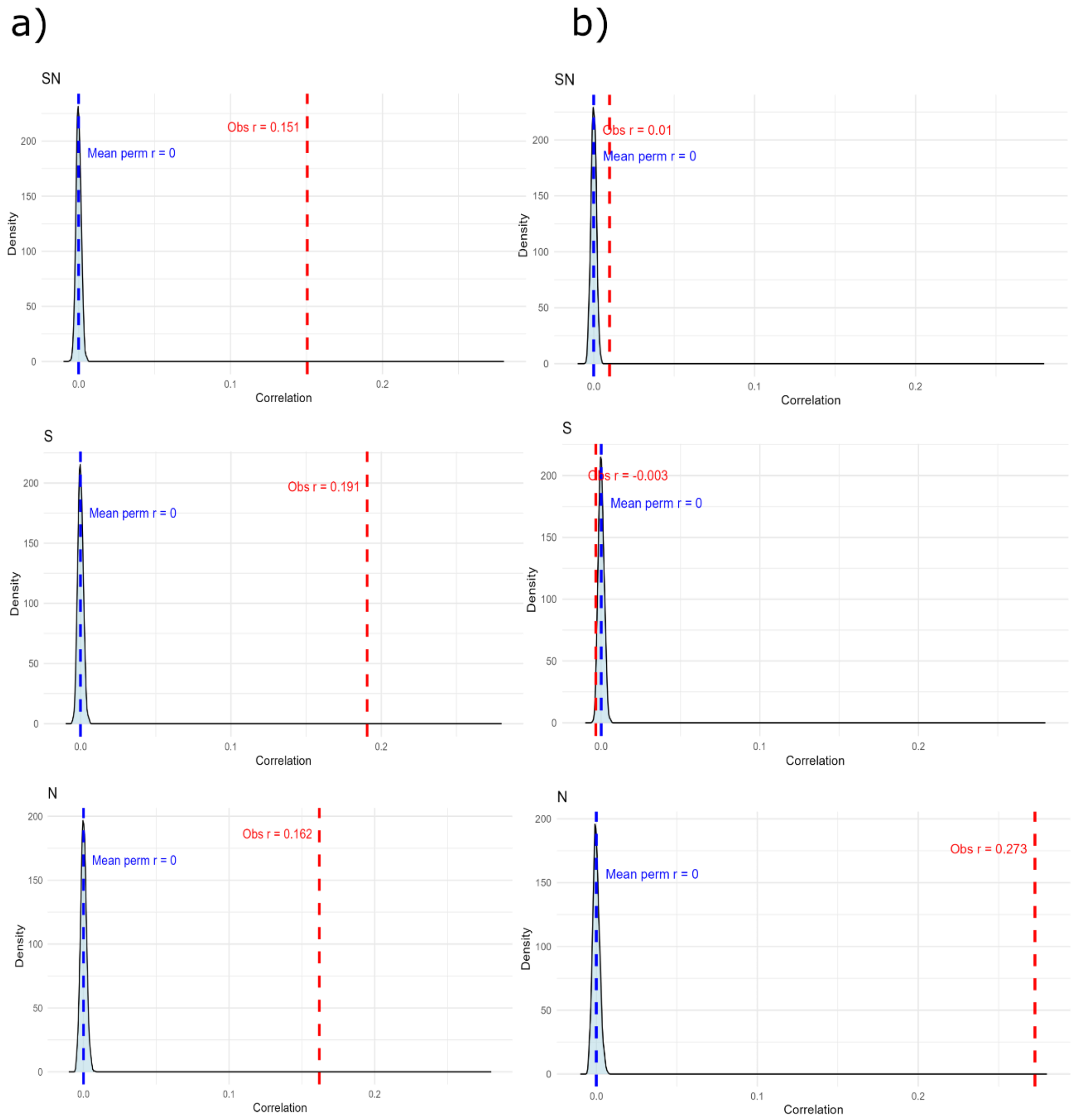


Figure S1.7 Permutation density plots of Pearson correlation coefficients between shared outliers in both nSL and Λ selection scans computed using (a) 52 and (b) 117 SNP window size in three population datasets: SN (top row), S (middle row), and N (bottom row). In each panel, the light blue shaded area represents the null distribution of Spearman's correlation coefficients calculated from 1000 permutations of the data; the blue dashed line marks the mean correlation coefficients for the permuted datasets ("Mean perm r"), and the red dashed line indicates the observed correlation ("Obs r").

#### 1.8 Permutation test to assess whether outlier windows form larger clusters within inversions than in collinear parts of the genome

To assess whether outliers tend to form larger clusters within inversions compared to collinear regions, we calculated the mean cluster size separately for inverted (region = 1) and collinear (region = 0) genomic windows. The observed difference in mean cluster size (inversion – collinear, in the table below "Difference mean cluster size (obs)") was then compared to a null distribution generated by randomly permuting the genomic region labels 1000 times, while preserving the outlier information column. For each permutation, we recalculated the mean cluster size within each genomic region type and computed the difference between them (in the table "Difference mean cluster size (perm)"). In all cases, observed cluster sizes within inversions were significantly larger than those in collinear regions.

Table S1.8. Mean difference in cluster size (inversion – collinear) observed (obs) and under permutation (perm) for outlier windows in Λ and nSL selection scans across three datasets (SN, S, N), using 52 and 117 SNP window sizes. Asterisks indicate statistically significant results ($p < 0.001$).

| Dataset | Window size | Statistic | Difference mean cluster size (obs) | Difference mean cluster size (perm) | p-value |
|---|---|---|---|---|---|
| SN | 52 | Λ | 53.38 | -33.92 | <0.001* |
| SN | 52 | nSL | 0.36 | -0.64 | <0.001* |
| SN | 117 | Λ | 301.15 | -106.75 | <0.001* |
| SN | 117 | nSL | 1.07 | -1.97 | <0.001* |
| S | 52 | Λ | 83.27 | -46.67 | <0.001* |
| S | 52 | nSL | 0.41 | -0.68 | <0.001* |
| S | 117 | Λ | 181.11 | -63.44 | <0.001* |
| S | 117 | nSL | 1.73 | -2.04 | <0.001* |
| N | 52 | Λ | 106.42 | -38.84 | <0.001* |
| N | 52 | nSL | 2.02 | -1.16 | <0.001* |
| N | 117 | Λ | 230.70 | -84.34 | <0.001* |
| N | 117 | nSL | 6.31 | -3.26 | <0.001* |

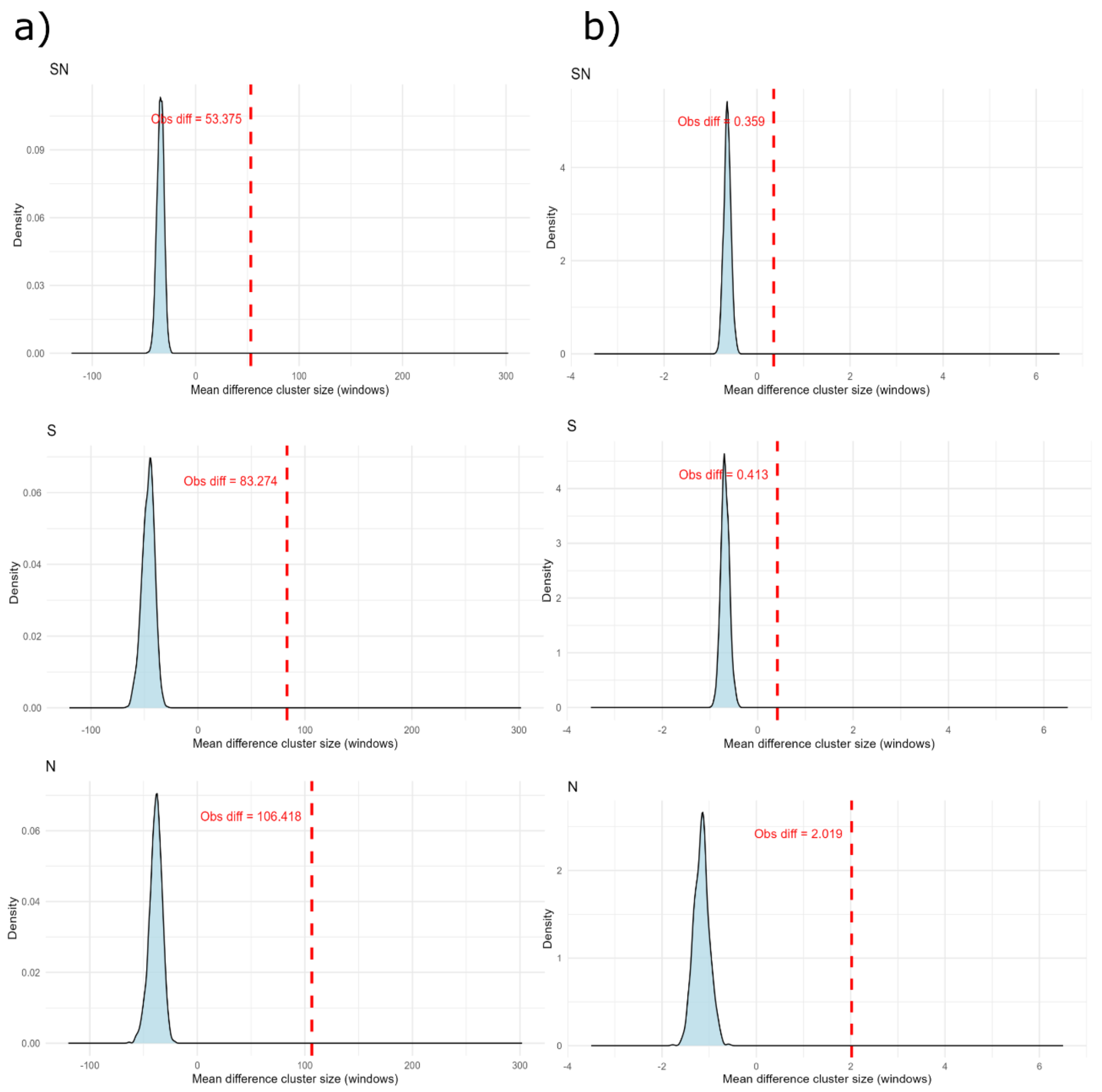


Figure S1.8.I Permutation density plots of mean difference in cluster sizes between inverted and collinear regions for both selection scans (a) Λ and (b) nSL using 52 SNP window size for the three populations datasets SN (top row), S (middle row) and N (bottom row). Each panel shows the distribution of mean cluster sizes calculated from 1000 permutations (light blue). The red dashed line indicates the observed mean difference in cluster size in windows ("Obs diff", inversion – collinear).

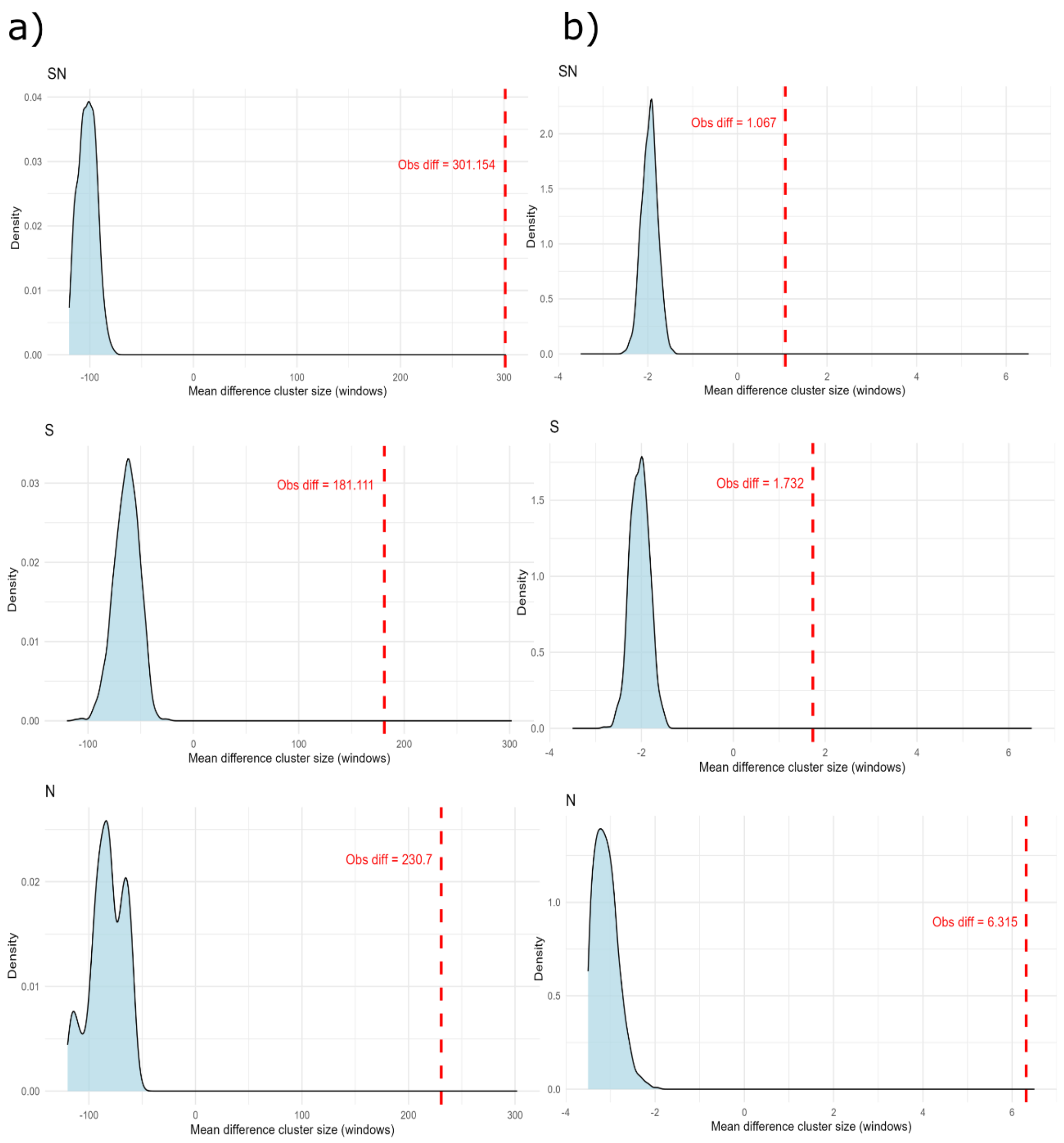


Figure S1.8.II Permutation density plots of mean difference in cluster sizes between inverted and collinear regions for both selection scans (a) Λ and (b) nSL using 117 SNP window size for the three populations datasets SN (top row), S (middle row) and N (bottom row). Each panel shows the distribution of mean cluster sizes calculated from 1000 permutations (light blue). The red dashed line indicates the observed mean difference in cluster size in windows ("Obs diff", inversion – collinear).

2. <u>Tables</u>

Table S1. *Ips typographus* sampling information. ‘Group’ indicates in what part of the species’ distribution area beetles were collected: northern (N), southern (S) and Poland (P). ‘N’ indicates the number of individuals collected per population.

| Population ID | N | Population name | Latitude | Longitude | Country | Group |
|---|---|---|---|---|---|---|
| ABE | 13 | Abetone | 44.148 | 10.663 | Italy | S |
| KAL | 15 | Kals | 47.012 | 12.646 | Austria | S |
| LIN | 14 | Linz | 48.092 | 13.874 | Austria | S |
| BAW | 11 | Bavarian Forest | 48.960 | 13.395 | Germany | S |
| TRE | 13 | Trebic | 49.212 | 15.879 | Czech | S |
| STE | 14 | Steigerwald | 49.622 | 10.263 | Germany | S |
| BIL | 14 | Bilkovice | 49.761 | 14.848 | Czech | S |
| SLA | 15 | Slanic | 45.250 | 25.925 | Romania | S |
| MOI | 14 | Moinesti | 46.510 | 26.449 | Romania | S |
| CAL | 15 | Calimani | 47.163 | 25.221 | Romania | S |
| ROZ | 14 | Roztocze | 50.508 | 22.786 | Poland | P |
| GOS | 14 | Goscinno | 54.047 | 15.657 | Poland | P |
| LUB | 14 | Lubaszki | 54.057 | 17.556 | Poland | P |
| BOR | 12 | Borki | 54.090 | 21.912 | Poland | P |
| TON | 13 | Tönnersjö | 56.643 | 13.070 | Sweden | N |
| ASA | 14 | Asa | 57.165 | 14.783 | Sweden | N |
| AAS | 13 | As | 59.667 | 10.793 | Norway | N |
| SIL | 14 | Siljanfors | 60.757 | 14.066 | Sweden | N |
| LAN | 13 | Länsi | 61.723 | 23.633 | Finland | N |
| EFI | 13 | Eastern Finland | 62.492 | 30.010 | Finland | N |
| STJ | 13 | Stjordal | 63.469 | 10.918 | Norway | N |
| SVA | 13 | Svartberget | 64.236 | 19.570 | Sweden | N |
| MEL | 14 | Mellakoski | 66.399 | 24.440 | Finland | N |

Table S2. Number of inversions that contained at least one outlier window in different datasets out of the total of 24 inversions (SN, S and N, both for Λ and nSL selection scans and 52 SNP window size).

| | | SN | S | N |
|---|---|---|---|---|
| Number of inversions with outliers | Λ | 4/24 | 6/24 | 3/24 |
| | nSL | 17/24 | 17/24 | 12/24 |

Table S3. Information on the longest outlier clusters identified in different dataset (SN, S, N) in Λ and nSL selection scans using 52 SNP window size. A cluster is considered when we find two or more consecutive outlier windows in a dataset. The table lists the genomic regions identified as the longest cluster of outliers in each dataset, indicates if those clusters are in collinear or inverted part of the genome, and indicates their respective coordinates (start - end) and size (in Mb).

| Dataset | Statistic | Contig | Start (Mb) | End (Mb) | Size (Mb) | Region |
|---|---|---|---|---|---|---|
| SN | Λ | 3 | 12.315 | 12.632 | 0.317 | collinear |
| SN | nSL | 5 | 5.969 | 5.985 | 0.016 | inverted |
| SNcol | Λ | 3 | 12.315 | 12.632 | 0.317 | collinear |
| SNcol | nSL | 3 | 12.544 | 12.564 | 0.020 | collinear |
| S | Λ | 16 | 0.830 | 1.154 | 0.324 | inverted |
| S | nSL | 5 | 5.691 | 5.737 | 0.046 | inverted |
| Scol | Λ | 3 | 12.337 | 12.657 | 0.320 | collinear |
| Scol | nSL | 3 | 10.224 | 10.254 | 0.030 | collinear |
| N | Λ | 5 | 4.697 | 4.950 | 0.253 | inverted |
| N | nSL | 5 | 4.863 | 4.899 | 0.036 | inverted |
| Ncol | Λ | 3 | 10.565 | 10.934 | 0.369 | collinear |
| Ncol | nSL | 3 | 10.224 | 10.262 | 0.038 | collinear |

Table S4. The comparison between Λ and nSL selection scans using different datasets and a 117 SNP window size. Calculations from nSL analysis are based on the absolute values of the statistic. Genomic region: part of the genome that is included in each dataset. Inversion-associated outliers: the percentage of outlier windows located within inversion regions, calculated from the total number of outliers. Collinear-region outliers: percentage of outliers located outside inversion regions calculated from the total number of outliers. Shared outliers: percentage of outliers that are shared between Λ and nSL results. Shared outliers in inversions: the percentage of outliers shared between Λ and nSL results that are in inversion regions. Shared outliers in collinear: the percentage of outliers shared between Λ and nSL results that are in the collinear genome. Information longest cluster: state in what contig we find the longest cluster and if it is inverted or collinear region. The notations (inv.) and (col.) refers to the genomic regions the cluster is located - inverted or collinear part respectively.

| Dataset | Genomic region | Statistic | Inversion associated outliers | Collinear region outliers | Shared outliers | Shared outliers in inversions | Shared outliers in collinear | Information longest cluster |
|---|---|---|---|---|---|---|---|---|
| SN | Inversions + collinear | Λ | 100% | 0% | 2% | 2% | 0% | Contig7 (inv.) |
| | | nSL | 84% | 16% | | | | Contig5 (inv.) |
| SNcol | Only collinear | Λ | | 100% | 21% | | 21% | Contig3 (col.) |
| | | nSL | | 100% | | | | Contig17 (col.) |
| S | Inversions + collinear | Λ | 100% | 0% | 1% | 1% | 0% | Contig7 (inv.) |
| | | nSL | 87% | 13% | | | | Contig5 (inv.) |
| Scol | Only collinear | Λ | | 100% | 15% | | 15% | Contig3 (col.) |
| | | nSL | | 100% | | | | Contig2 (col.) |
| N | Inversions + collinear | Λ | 100% | 0% | 28% | 28% | 0% | Contig5 (inv.) |
| | | nSL | 93% | 7% | | | | Contig5 (inv.) |

| | | | | | | | | |
|---|---|---|---|---|---|---|---|---|
| Ncol | Only collinear | Λ<br>nSL | | 100%<br>100% | 21% | | 21% | Contig3 (col.)<br>Contig3 (col.) |

Table S5. Percentage of outliers per inversion in different datasets including all individuals (SNInv), major allele homozygotes (MJAInv), and minor allele homozygotes (MNAInv). Those datasets include all collinear regions and only the inverted part that is analysed, the rest of the inversions were excluded in each analysis. Upper panel: results for nSL; lower panel: results for Λ. Orange colour means that the percentage of the signal is lower compared with the dataset with all individuals; pink means that the signal increases; grey means that the signals do not change comparing with SNInv dataset.

| | Signals in nSL | | | | | | | | | | | |
|---|---|---|---|---|---|---|---|---|---|---|---|---|
| | Inv2 | Inv3 | Inv5 | Inv6 | Inv7.1 | Inv7.2 | Inv10 | Inv12 | Inv13 | Inv14.1 | Inv14.2 | Inv14.3 |
| SNInv | 42% | 0.04% | 51% | 1.2% | 13% | 39% | 0% | 0% | 7% | 16% | 4% | 0.3% |
| MJAInv | 34% | 0% | 14% | 1.4% | 4% | 6% | 0% | 0% | 0.3% | 12% | 0.4% | 0% |
| MNAInv | 4% | 0% | 62% | 4% | 0% | 8% | 0% | 0% | 7% | 0.3% | 0% | 0.1% |
| | Inv14.4 | Inv14.5 | Inv15 | Inv16 | Inv17 | Inv18 | Inv22.1 | Inv22.2 | Inv22.3 | Inv22.4 | Inv23 | Inv26 |
| SNInv | 0% | 0% | 2% | 52% | 0.3% | 8% | 0% | 0% | 0% | 0% | 0% | 0.04% |
| MJAInv | 0% | 0.1% | 0% | 1% | 0% | 0.1% | 0.1% | 0% | 0.5% | 0% | 2% | 0% |
| MNAInv | 0% | 0% | 0% | 20% | 0% | 0.4% | 0% | 0.1% | 4% | 0.3% | 6% | 0.1% |
| | Signals in Λ | | | | | | | | | | | |
| | Inv2 | Inv3 | Inv5 | Inv6 | Inv7.1 | Inv7.2 | Inv10 | Inv12 | Inv13 | Inv14.1 | Inv14.2 | Inv14.3 |
| SNInv | 3% | 0% | 24% | 0% | 0% | 0% | 0% | 0% | 0% | 7% | 0% | 0% |
| MJAInv | 0.5% | 0% | 0.5% | 3% | 2% | 0% | 0% | 0% | 0% | 0.2% | 0% | 0% |
| MNAInv | 43% | 0% | 100% | 3% | 0% | 100% | 0% | 0% | 0% | 0% | 0% | 0% |
| | Inv14.4 | Inv14.5 | Inv15 | Inv16 | Inv17 | Inv18 | Inv22.1 | Inv22.2 | Inv22.3 | Inv22.4 | Inv23 | Inv26 |
| SNInv | 0% | 0% | 0% | 0% | 0% | 0% | 0% | 0% | 0% | 0% | 5% | 0% |

| | | | | | | | | | | | | |
|---|---|---|---|---|---|---|---|---|---|---|---|---|
| MJAInv | 0% | 0% | 0% | 0% | 0% | 0% | 0% | 0% | 0% | 0% | 7% | 0% |
| MNAInv | 0% | 0% | 0% | 30% | 0% | 0% | 0% | 0% | 0% | 0% | 0% | 0% |

Table S6. Gene Ontology (GO) terms significantly enriched in regions identified as selection outliers in *Ips typographus*. Fold enrichment was calculated as the ratio between the number of significant genes associated with a GO term and the number expected under a random distribution (Significant / Expected). Asterisks indicate levels of statistical significance based on eFisher p-values: $p < 0.05$*, $p < 0.01$**, and $p < 0.001$***. Datasets refers to signals that survived at least in one homozygote haplotype in Λ (Λ homozygotes) and nSL (nSL homozygotes), and in SN, S and N datasets, or just in one of them (S specific, N specific).

| GO ID | Term | Fold enrichment | eFisher p-value | Dataset |
|---|---|---|---|---|
| **GO:0071555** | Cell wall organization | 10.21 | 0.0156* | Λ homozygotes, nSL homozygotes |
| **GO:0030054** | Cell junction | 7.14 | 0.00042*** | Λ homozygotes |
| **GO:0005506** | Iron ion binding | 4.43 | 0.0025** | Λ homozygotes, nSL homozygotes |
| **GO:0004553** | Hydrolase activity | 2.89 | 0.0034** | Λ homozygotes |
| **GO:0005886** | Plasma membrane | 3.31 | 0.0011** | Λ homozygotes |
| **GO:0016705** | Oxidoreductase activity | 3.59 | 0.0058** | Λ homozygotes |
| **GO:0004497** | Monooxygenase activity | 4.09 | 0.0216* | Λ homozygotes, nSL homozygotes |
| **GO:0005230** | Extracellular ligand-gated monoatomic ion channel activity | 7.33 | 0.0173* | Λ homozygotes, nSL homozygotes |
| **GO:0020037** | Heme binding | 3.35 | 0.0080** | Λ homozygotes |
| **GO:0038023** | Signalling receptor activity | 3.15 | 0.0060** | nSL homozygotes |
| **GO:0050896** | Response to stimulus | 1.87 | 0.0038** | nSL homozygotes |
| **GO:0007034** | Vacuolar transport | 8.82 | 0.0041** | nSL homozygotes |
| **GO:0003964** | RNA-directed DNA polymerase activity | 2.92 | 0.0087** | SN, S, N |
| **GO:0006278** | RNA-templated DNA biosynthetic process | 2.58 | 0.0186* | SN, S, N |
| **GO:0004190** | Aspartic-type endopeptidase activity | 5.76 | 0.0104* | SN, S |
| **GO:0004519** | Endonuclease activity | 2.75 | 0.0147* | SN, S |
| **GO:0030246** | Carbohydrate binding | 7.33 | 0.0311* | SN, S |
| **GO:0043170** | Macromolecule metabolic process | 1.38 | 0.0331* | SN, S |

| | | | | |
|---|---|---|---|---|
| **GO:0006807** | Nitrogen compound metabolic process | 1.32 | 0.0452* | SN, S |
| **GO:1902531** | Regulation of intracellular signal | 11.51 | 0.0124* | SN, N |
| **GO:0007264** | Small GTPase mediated signal | 9.17 | 0.0197* | SN, N |
| **GO:0003779** | Actin binding | 6.51 | 0.0377* | SN, N |
| **GO:0004803** | Transposase activity | 15.38 | 0.0067** | S specific |
| **GO:0016301** | Kinase activity | 4.60 | 0.0098** | N specific |

3. <u>Figures</u>

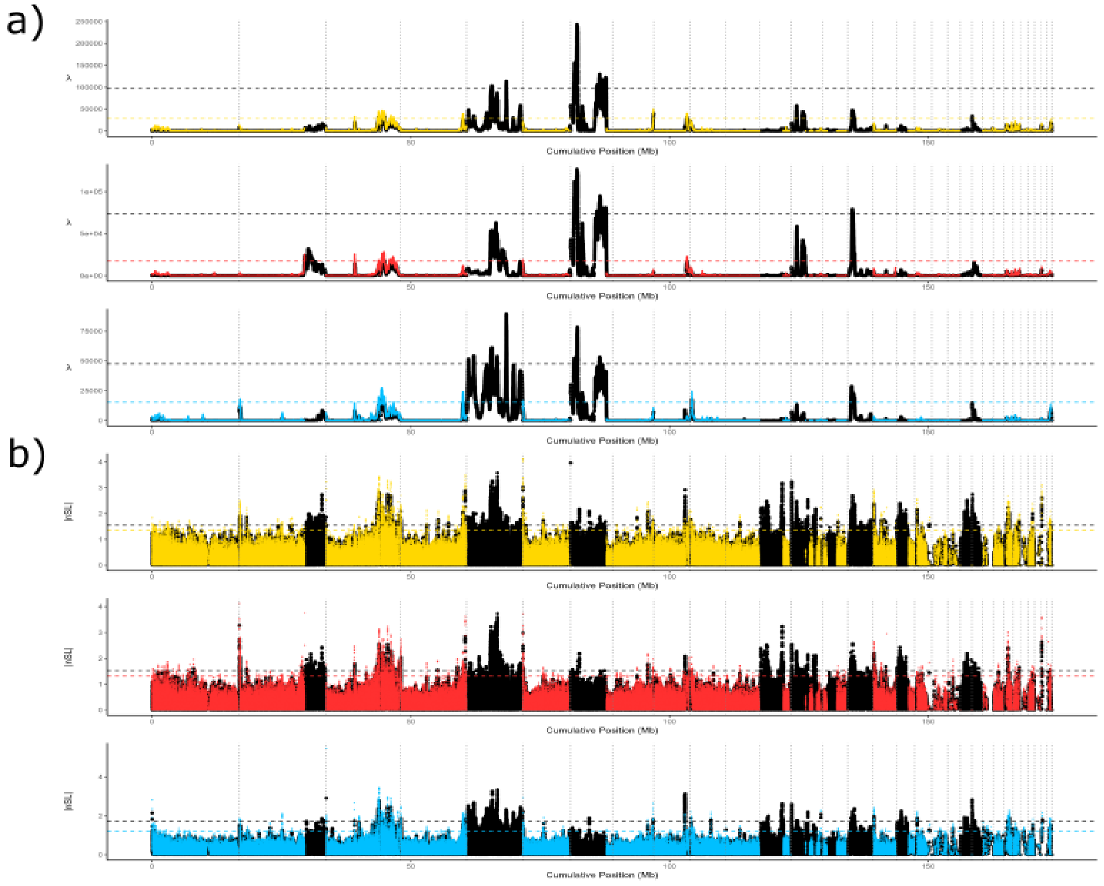


<u>Figure S1</u>. Results of the selection scans for Λ (a) and nSL (absolute values) (b) with 117 SNPs window sizes and the SN dataset (top panel), the S dataset (middle panel), and the N dataset (bottom panel). The whole genome (both collinear and inverted regions included) results are shown in black. The same analysis was performed excluding inversion regions, and the results are overlaid on the whole-genome results in different colors for the different datasets: SN (yellow), S (red), and N (blue). The dashed lines indicate the outlier threshold (1% of all values). In all plots, gray dotted vertical lines represent the end of each contig.

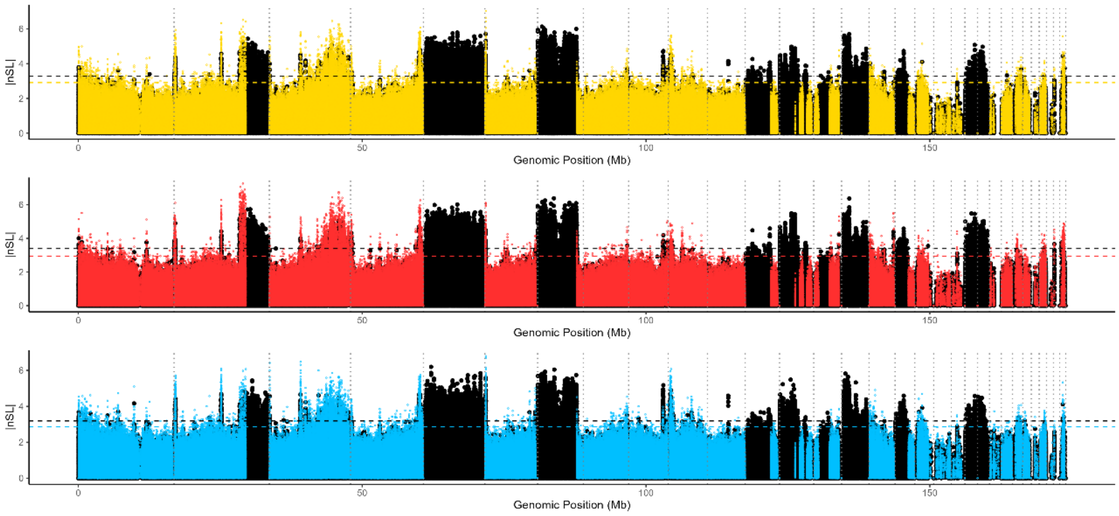


Figure S2. Selection scans results for normalised values of nSL (absolute values) without windowing. Top panel: SN dataset; middle panel: S dataset; bottom panel: N dataset. The whole-genome results (both collinear and inverted regions included) are shown in black, with the same analysis excluding inversion regions overlaid on the whole-genome results in different colors for the different datasets: SN (yellow), S (red), and N (blue). The dashed lines indicate outliers threshold (1% of all the values). In all plots, grey dotted vertical bars represent the end of each contig.

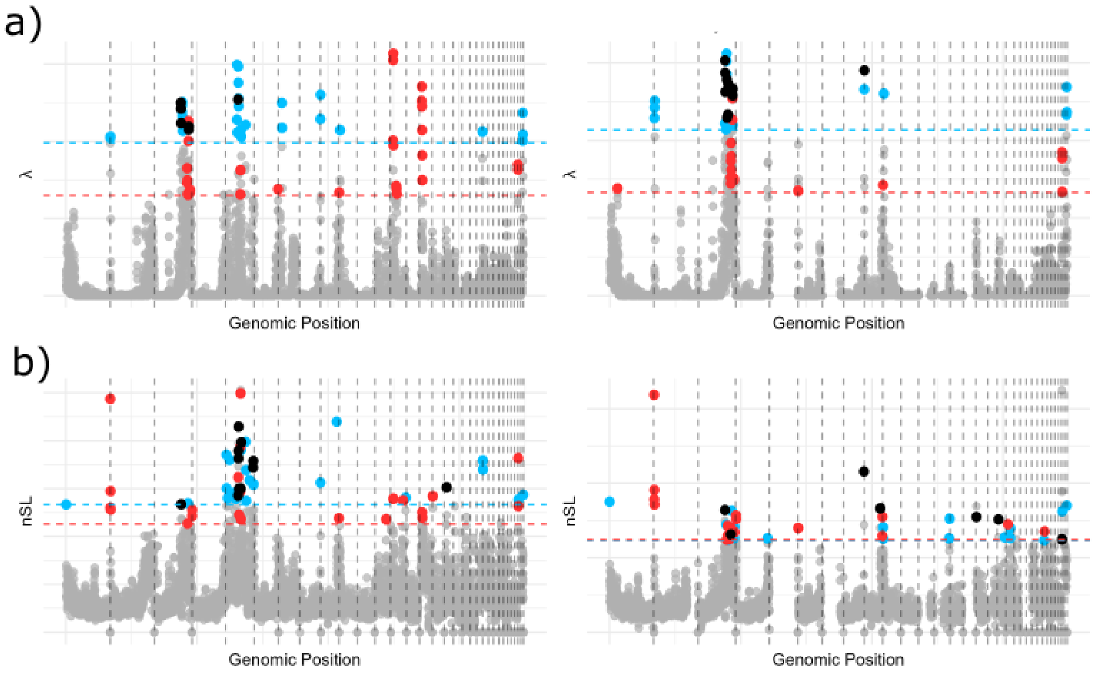


Figure S3. Comparison of the identified outliers between S and N datasets in (a) Λ and (b) nSL (absolute values) selection scans using 50kb windows. Left plots include datasets with inversions and collinear parts, and right plots include only collinear parts. Outliers: appearing in both south and north datasets (black), only in north (blue), only in south (red). Vertical lines define the end of the contigs.

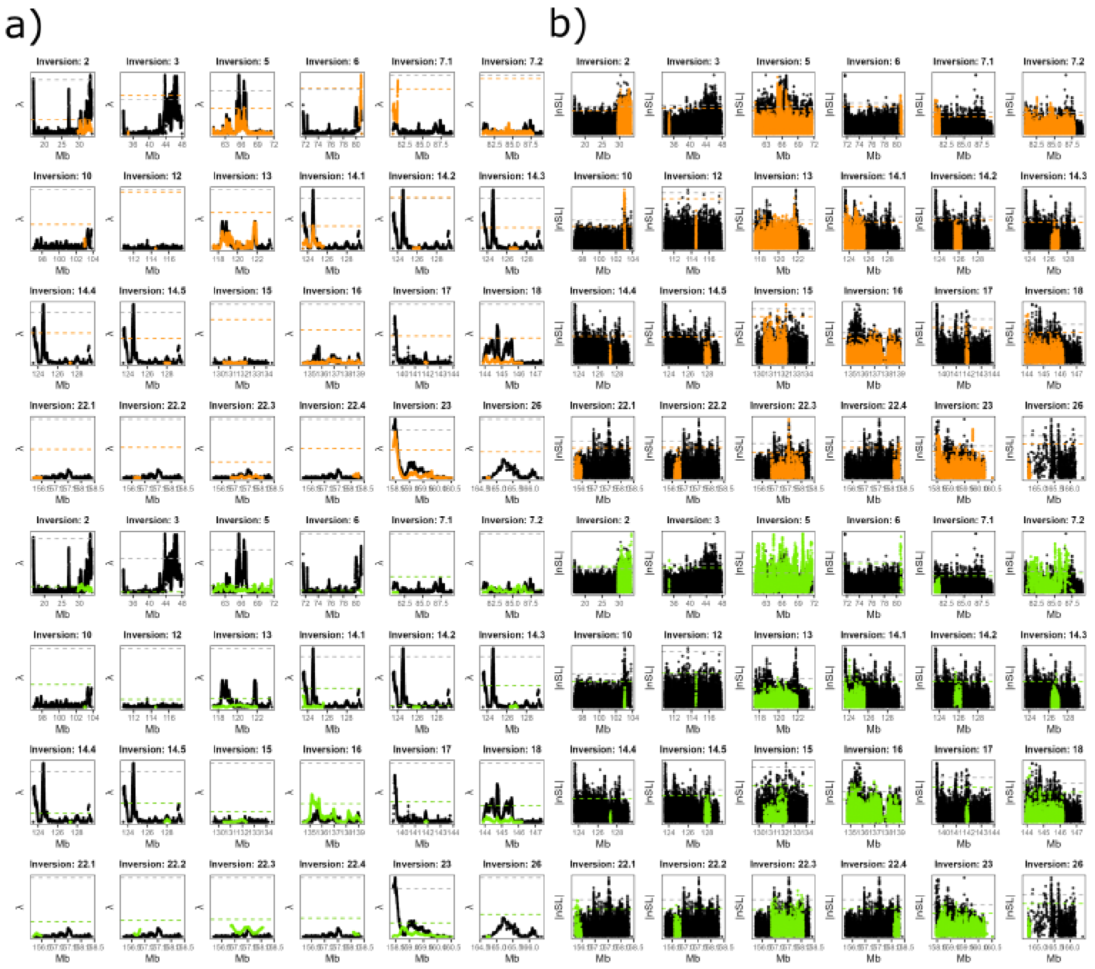


Figure S4. Selection signals in the inversions in the context of contigs (each panel shows a contig with inversion). (a) Λ results. Results obtained using datasets that included all possible genotypes are shown in black and results obtained using datasets That included only inversion homozygotes are shown in orange (major allele homozygotes) or green (minor allele homozygotes). The threshold is set at the 1% values of Λ and normalized nSL (absolute values) respectively to identify significant selection signals (dashed lines). (b) nSL results follow the same layout and interpretation as in (a).
